# Data-Driven Modelling of Gene Expression States in Breast Cancer and their Prediction from Routine Whole Slide Images

**DOI:** 10.1101/2023.04.14.536756

**Authors:** Muhammad Dawood, Mark Eastwood, Mostafa Jahanifar, Lawrence Young, Asa Ben-Hur, Kim Branson, Louise Jones, Nasir Rajpoot, Fayyaz ul Amir Afsar Minhas

## Abstract

Identification of gene expression state of a cancer patient from routine pathology imaging and characterization of its phenotypic effects have significant clinical and therapeutic implications. However, prediction of expression of individual genes from whole slide images (WSIs) is challenging due to co-dependent or correlated expression of multiple genes. Here, we use a purely data-driven approach to first identify groups of genes with co-dependent expression and then predict their status from (WSIs) using a bespoke graph neural network. These gene groups allow us to capture the gene expression state of a patient with a small number of binary variables that are biologically meaningful and carry histopathological insights for clinically and therapeutic use cases. Prediction of gene expression state based on these gene groups allows associating histological phenotypes (cellular composition, mitotic counts, grading, etc.) with underlying gene expression patterns and opens avenues for gaining significant biological insights from routine pathology imaging directly.

**Graphical Abstract:** 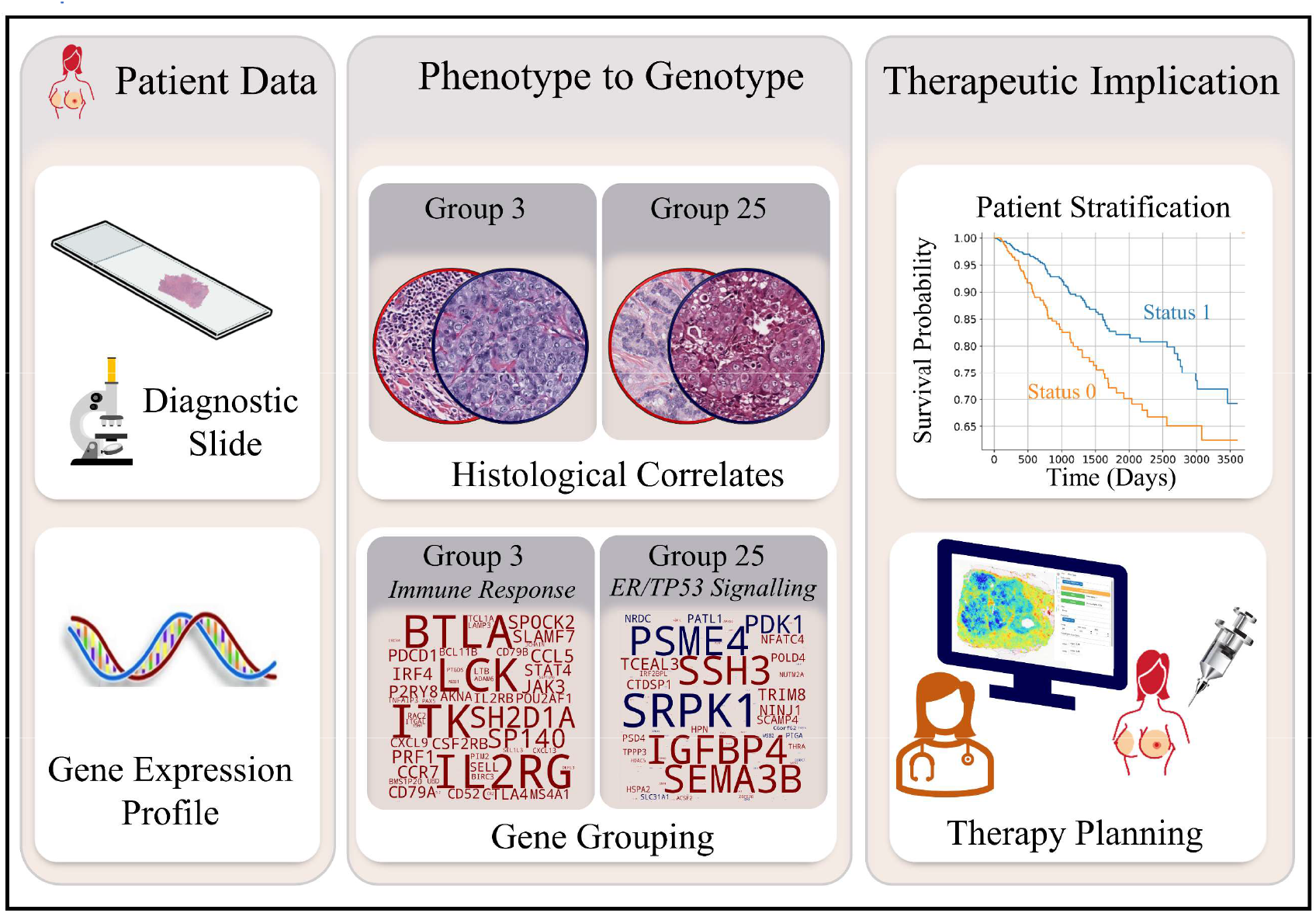

**Highlights:**

- Data-driven discovery of co-expressing gene groups in breast caner
- Histological imaging based prediction of gene groups via deep learning
- Identification of phenotypic correlates of gene-expression in histological imaging
- Clinical and therapeutic impact of gene groups and their visual patterns identified

## 1 Introduction

Cancer is a clonal disease in which genetic alterations directly or indirectly alter gene expression, biological pathways, and proteins activity leading to phenotypic changes in the spatial organization of the tumor microenvironment (TME) [1]. Consequently, associating histological and molecular patterns is crucial for understanding disease mechanism and clinical decision-making [2]. Like other cancers, breast tumors also exhibit heterogeneity at both morphological and molecular levels and are divided into several histological and molecular subtypes. During histopathology examination, a tumor section stained with Hematoxylin and Eosin (H&E) is visually examined for features such as mitotic counts, nuclear pleomorphism, epithelial tubule formation, necrosis and tumor-infiltrating lymphocytes, etc., to develop a spatially-informed histological profile of the disease. Similarly, gene expression analysis based on molecular tests such as PAM50 [3], [4], Oncotype-Dx [5] and Mammaprint [6] can also be used for patient subtyping. Gene expression profiling based on such limited gene assays or from Bulk RNA-Seq [7] and single-cell RNA-sequencing (scRNA-seq) [8], [9] plays a key role in understanding the genetic basis of cancer and discovery of novel therapeutic targets. However, such technologies are unable to capture spatial heterogeneity in the expression profile of genes across a tumor section. Spatial profiling of a tumor transcriptome is typically achieved using Spatially resolved Transcriptomics (SpTx) technologies [10]. However, such technologies are generally costly and offer low resolution in terms of spatial details or genes [11], [12]. Consequently, there is a need for cross-linking gene expression and spatial histological imaging profiles to gain a more in-depth understanding of latent factors associated with the disease.

In an attempt to achieve this goal, recent advancements in deep learning for computational pathology have demonstrated that prediction of expression profiles of genes is possible from whole slide images (WSIs) of H&E stained tissue sections [13]–[15]. For example, Schmauch et al. proposed a deep learning method called HE2RNA for predicting gene expression profiles from WSIs. Similarly, Wang et al. proposed a deep learning method for predicting the expression profile of 17,695 genes from WSIs [16]. For each of the 17,695 genes, the authors have tiled the WSIs into patches and then trained and optimized an Inception V3 for predicting tile-level and WSI-level expression. Most recently, an attention-based called tRNAsformer has been proposed for predicting the expression level of the individual gene from WSIs in kidney cancer [17].

The vast majority of image-based RNA-Seq expression prediction methods focus on associating tissue morphology with the expression level of *individual* genes [15]–[17]. This is typically done by designing a machine learning pipeline in which the input is a WSI, and the output is the expression level of a single gene. However, due to the nature of the biological mechanisms underlying gene expression, genes usually show co-dependent or correlated expression. Consequently, it is, in general, not possible to associate the predicted expression of a single gene from the input WSI to that gene alone. Furthermore, an observed phenotypic effect cannot solely be pinpointed to the known function of a single gene as, typically, it will be a collective effect exhibited by the expression of functionally interrelated genes and a single gene may be associated with multiple functions [18]. Therefore, instead of predicting the phenotypic effect of a single gene from WSIs, it is more meaningful to predict the expression of groups of genes that act concomitantly and exhibit coherent patterns of expression across samples.

In contrast to existing research in this domain that focuses on prediction of expression level of individual genes from WSIs, in this work we first characterize the gene expression state of a patient in terms of a small number of binary latent factors or gene groups that are discovered in a purely data-driven manner. These can be viewed as overlapping groups of related genes whose expression shows significant inter-dependence across samples. The motivation behind such gene grouping is that, though co-expression is not causation, co-expressed genes show coordinated responses across a significant subgroup of patients hinting that these genes may be part of an underlying biological pathway, protein complexes or disease subtype [19]. We have shown that the discovered gene groups are clinically and pathologically relevant in terms of their association with survival, breast cancer receptor status, histopathological phenotypes, cancer driver genes mutations, biological pathways enrichment and underlying protein-protein interactions, and also therapeutic decision-making. We then propose a bespoke multi-output graph neural network-based computational pathology pipeline to predict the expression state of a patient in terms of these latent factors from their WSIs. This enables identification of spatial histological patterns associated with individual latent factors as well as the overall gene expression profile of a patient. Finally, we have shown that image-based predicted gene group statuses can be used as a latent representation for the prediction of several other downstream clinical tasks such as patient subtyping, and also driver gene alteration status and pathway alteration status.

## 2 Results

### 2.1 Analytic workflow

As shown in **Fig 1**, we performed gene expression analysis of the TCGA breast cancer (TCGA-BRCA) cohort (*n* = 1084) to identify 200 groups of genes such that the expression of genes in the same group is maximally statistically co-dependent. This allows us to capture the inter-dependence between expression profiles of different genes and represent the gene expression state of a given patient in the form of 200 binary variables each corresponding to a single group. To underscore the clinical, therapeutic, and biological significance of each gene group, we computed the association of patient gene group status with survival, enrichment for biological pathways and cancer hallmark processes, and also protein-protein and drug-protein interactions.

**Figure 1:**
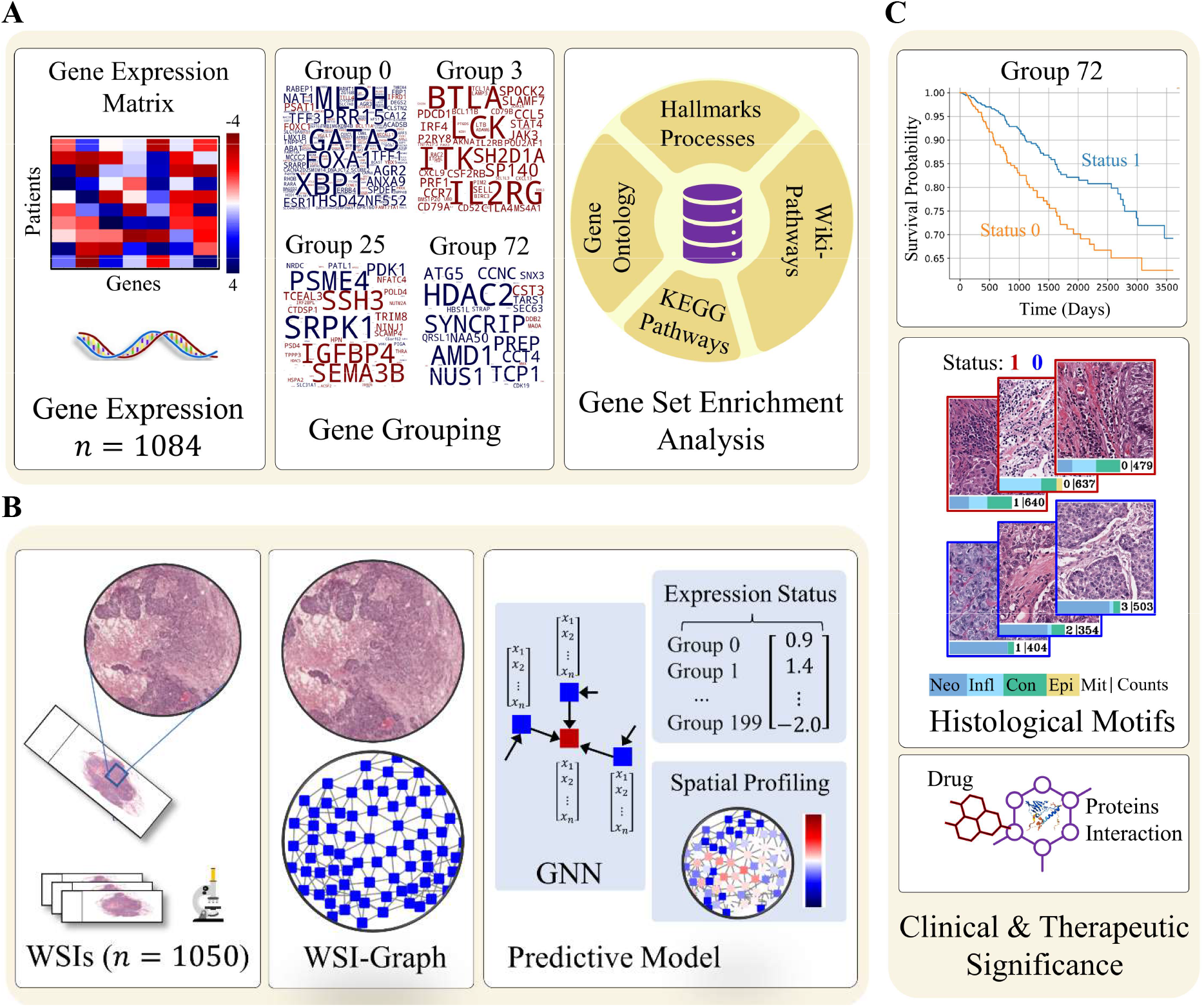
Analytic workflow for patient gene expression state prediction from whole slide images (WSIs). A) Workflow of data-driven discovery of gene groups and their pathological significance is shown. We first identified 200 binary latent factor or gene groups from the gene expression data in a data-driven manner. A gene group can be viewed as overlapping group of genes that exhibit coherent patterns of expression across sample. Word clouds demonstrating the gene composition of different gene groups. The color of the gene indicates whether its median expression across patients is high (red) or low (blue) when gene group status = 1. Afterward, we assessed the biological significance of the genes grouped in different gene groups through gene set enrichment analysis. B) The proposed 𝑆𝑙𝑖𝑑𝑒𝐺𝑟𝑎𝑝ℎ^oo^ pipeline for prediction of gene groups status from WSIs. We first construct graph representation of a WSI and then feed it into a Graph Neural Network (GNN) for predicting WSI-level and spatially resolved expression status of these 200 gene groups. C) Identification of clinically relevant gene groups in term of association with survival and their associated histological motifs. Histology image-based inference of personalized medication by analyzing protein-protein and drug-protein interaction of gene groups.

We then used our bespoke graph neural network-based pipeline that takes a WSI as input and predicts the binary status of 200 gene groups simultaneously in an end-to-end manner. This allows us to model the complete gene expression profile of a patient and identify histological imaging patterns associated with each gene group. Furthermore, the proposed approach allows spatially resolved cross-linking of discovered gene groups with visual information contained in the WSI. The interactive visualization portal for the proposed approach (called Histology Gene Groups Xplorer (HiGGsXplore)) is available at: (http://tiademos.dcs.warwick.ac.uk/bokeh_app?demo=HiGGsXplore)

### 2.2 Data-Driven discovery of Gene Groups based on co-dependent expression

To capture multivariate nonlinear relationships in gene expression patterns across patient samples, we employed Correlation Explanation (CorEx) on RNA-Seq data of the TCGA-BRCA cohort. CorEx can be used to model the underlying dependency structure of a dataset by identifying groups of random variables that in the context of this application can intuitively be viewed as a manifestation of underlying covarying patterns of gene expression profiles of different genes across patients. The input to CorEx is a 1084 × 5676 matrix where each row is the normalized gene expression score of 5,676 genes with high expression variance or mutation frequency for each of the 1,084 patients. For this data, CorEx identified 200 gene groups that can explain the co-dependence between gene expression patterns observed in the data without loss of information. This allows us to represent the gene expression state of each patient in terms of these 200 binary variables rather than the expression of all individual genes. As these gene expression groups are identified in a purely empirical manner from gene expression data, the expected impact of any human observation biases on the definition of these gene groups is minimal. Furthermore, a single gene can be associated with multiple gene groups which is desirable from a biological point of view as gene products often perform multiple roles within a cell and can be part of multiple interaction networks [20].

The gene composition of a selected number of gene groups is shown as word clouds in **Fig 2A** and **SFig 1**. For example, the binary status of Gene Group 0 (G0) is defined primarily based on the expression patterns of a set of genes (*MLPH, GATA3, XBP1, FOXA1, TFF3, ESR1,* etc.). The exhaustive list of genes grouped in all 200 gene groups is provided in supplementary data. **Fig 2B** illustrates the underlying co-dependent expression of genes grouped in a selected gene group along with their group status. The heatmaps clearly show that the expression level of genes in Gene Group 3 (G3) and Gene Group 25 (G25) are significantly co-dependent across patients. For instance, for patients with G3 = 1, the expression level of *ITK, IL2*, *PDCD1* or *PD1, ITGAL, PDCD1LG2* or *PD-L2*, and several other genes are high, whereas, for patients with G3 = 0, these genes show under-expression as evident from the figure. For G25, a consistent trend in gene expression can be seen between status = 0 and 1 patients. For example, for patients with G25 status = 1, *MYC, CHEK1, PSME4, YES1, NRAS*, *TP53*, and several other genes show high expression levels, whereas, *IGFBP4, TCEAL3*, *RORC, RETSAT,* and others show low expression. Conversely, for patients with G25 = 0, the expression patterns of these genes are reversed.

**Figure 2:**
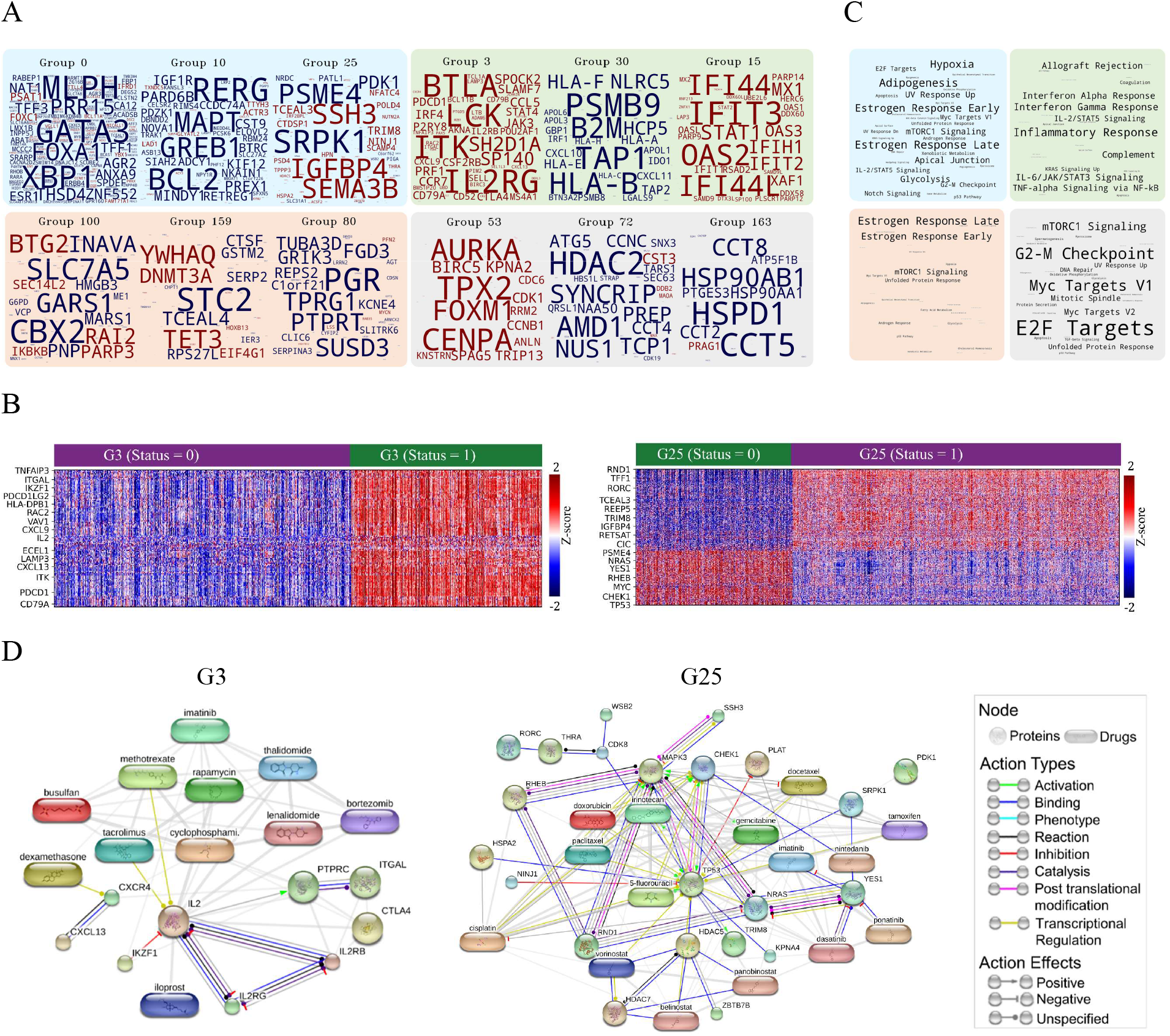
Data Driven Discovery of Gene Groups, their biological and therapeutic significance. A) Word clouds demonstrating the gene composition of different gene groups. The color of the gene indicates whether its median expression across patients is high (red) or low (blue) when gene group status = 1. The font size of gene within a group is proportional to the amount of information that the gene status provides about a particular gene. B) Gene expression profile and group status of genes (one per row) for all patients (one per column) in Gene Group 3 (G3) and Gene Group 25 (G25) are shown. C) Enriched terms for hallmark processes in similar gene groups (note color in A) are shown, with font sizes proportional to the number of gene groups that show enrichment for a certain process. D) Protein-protein and protein-drug interaction of selected genes in G3 (left plot) and G25 (right plot) are shown. Nodes shown in circles represent proteins, while the rounded rectangle shapes represent drugs. The edges between nodes show different types of interaction and potential therapeutic targeting.

This key result lends support to the motivation of this work, i.e., the expression level of multiple genes is significantly and consistently inter-dependent and the overall gene expression state of a patient can be characterized by a small number of latent factors. It also highlights the fact that it is not possible to disentangle the expression status of individual genes and consequently associate an observed phenotype, say in a WSI, with the status of a single gene. We next investigated the pathological significance of these gene groups and analyze their predictability from WSIs.

### 2.3 Pathological Significance of Gene Groups

Here we discuss the clinicopathological significance of gene groups to understand the implications of these latent factors for clinical decision-making before analyzing their predictability from imaging.

#### 2.3.1 Association of Gene Groups with Cancer Hallmarks and Biological Pathways

Through Gene Set Enrichment Analysis (GSEA) we found genes from several gene groups associated with known cancer hallmark processes and biological pathways. In **Fig 2C** we show the enriched terms for cancer hallmark processes in selected gene groups. For example, genes in Gene Group 0, 10 and 25 show enrichment for Estrogen early and late response, KRAS and mTORC1 signalling, Unfolded Protein Response (UPR), p53 pathway and several other hallmark processes. Similarly, we found genes from Gene Group 3, 15 and 30 associated with Inflammatory response, Interferon Alpha and Gamma response, and several other cancer hallmark processes. Additionally, we found genes from several gene groups associated with several cancer hallmark processes (Epithelial-Mesenchymal Transition (EMT), Myc targets V1 and V2, Mitotic spindle, DNA repair, KRAS up and down signalling, etc.) as shown in **SFig 2**.

Apart from cancer hallmark processes, several a number of gene groups has shown enrichment enriched for several biological processes (e.g. T-cell receptor signalling, MAPK cascade, negative regulation of programmed cell death, etc.,) KEGG pathways (e.g. *PD-L1* expression and *PD-1* checkpoint pathway, JAK-STAT and PI3K-Akt signalling pathway, Th1, Th2 and Th17 cell differentiation, etc.,) and WikiPathways (e.g. DNA damage response, Inflammatory response, B Cell receptor signalling, etc.,) as can be seen in **SFig 3**, **SFig 4** and **SFig 5**. For example, G3 and several other gene groups have shown enrichment for *PD-L1* expression and *PD-1* checkpoint pathway in cancer which can be a guiding signal for therapeutic decision-making [21].

#### 2.3.2 Gene Groups capture clinically important protein-protein and protein-drug interactions

We analyzed the protein-protein interaction (PPI) and protein-drug interaction (PDI) of genes in several gene groups with the end goal of identifying which groups involve proteins that can be targeted with known drugs so that the gene group status can be used as a potential indicator to guide therapeutic decision making. **Fig 2D** shows the PPI and PDI of a selected number of genes from G3 and G25. Regarding G3, interaction between *IL2, IL2RB* and *IL2RG* can be seen (left figure), which is expected as *IL2* regulates immunity by teaming up with *IL2RB* and *IL2RG* [22], [23]. Similarly, interaction of tacrolimus, an immunosuppressive and anti-inflammatory macrolide that targets the CD4+-cells can be seen with *IL2*. As these genes show high expression when G3 = 1, therefore patients with G3 = 1 can be considered a candidate for tacrolimus therapy. In reference to G25 (see right figure), TRIM8 a member of the tripartite motif-containing (TRIM) binding with TP53 can be seen, which has been shown to play a role in regulating TP53/p53- mediated pathway [24]. Similarly, interaction of *YES1*, a targetable oncogene can be seen with drugs such as dasatinib, ponatinib, nintedanib and imatinib. When G25 = 0, *YES1* shows high expression, therefore patients with G25 = 0 could be considered as potential candidates for dasatinib therapy [25]. Apart from this, interaction of *TP53* with several other proteins (*CHEK1, MAPK3, PLAT, NINJ1, HDA*C5, etc.) and drugs (tamoxifen, doxorubicin, paclitaxel, etc.) can be observed.

#### 2.3.3 Patient stratification into high and low risk using gene groups status

We found the binary status of several gene groups associated with overall survival (OS), disease-specific survival (DSS), and progression-free survival (PFS) of patients. **Fig 3A** shows the Kaplan-Meier (KM) survival curves (DSS, PFS and OS) illustrating patients’ stratification based on their gene group status. The KM curves indicate that patients can be stratified into high and low risk groups based on their G25 and G195 status with statistical significance (log-rank test FDR-corrected p-value > 0.05). Additionally, from the figure, patients with G3 = 1 have higher survival rates compared to those with G3 = 0 but the stratification is not statistically significant. Our analysis shows that the number of gene groups with statistically significant risk stratification (multiple-hypothesis corrected log-rank p-value < 0.05) is 25, 3 and 2 for DSS, OS and PFS, respectively as shown in **SFig 6**.

**Figure 3:**
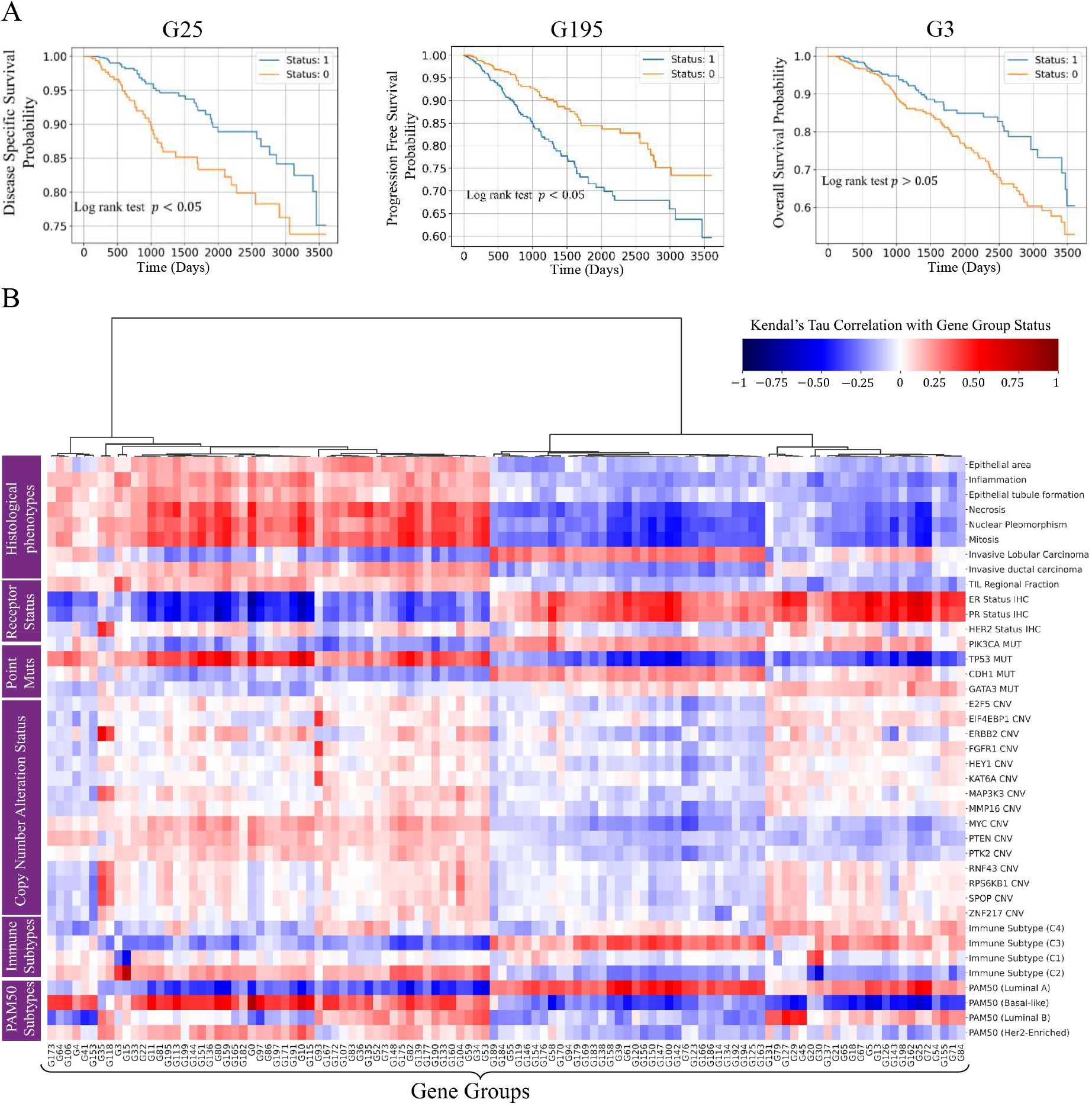
Clinical and pathlogical significane of gene groups binary status. A) Kaplan-Meier curve showing stratification of patient into high and low risk group based on G3 (left plot), G25 (middle plot) and Gene group 195 (G195) binary status. G25 and G195 status of a patient is associated with 10-year censored disease specific survival and progression free survival (log rank test FDR corrected p-value < 0.05). G3 status can stratify patient into high and low risk group but FDR corrected p-value is not significant. B) Association of gene groups with histological phenotypes, receptor status, genes point mutation status and copy number alteration status, and also immune and PAM50 molecular subtypes. Gene groups are shown along x-axis, and histological phenotypes and other clinical markers are shown along y-axis. Red and blue colors indicate the degree of association between gene groups status and a specific histopathological phenotype or clinical marker. Dark-red color shows strong positive correlation while strong negative correlation is shown using dark-blue color. (Abbreviations -CNV: Copy Number Variations, TIL: Tumor Infiltrating Lymphocytes)

#### 2.3.4 Association between Gene Groups and breast cancer receptor status

We found the status of several gene groups associated with ER, PR and Her2 status as can be seen in **Fig 3B**. For example, from the figure strong positive association of G25 status with ER (Kendall-tau correlation coefficient 𝜌_r_ = 0.68 and 𝑝 < 0.01) and PR (𝜌_r_ = 0.58 and 𝑝 < 0.01) status can be seen. This correlation was expected as G25 status is defined by IGFBP4 and other relevant genes whose overexpression has previously been found positively associated with ER and PR status [26]. Similarly, we found G35 and G118 status strongly positively associated with her2 status as evident from the figure.

#### 2.3.5 Association with PAM50 molecular subtypes and immune subtypes

We found the status of several gene groups associated with PAM50 molecular subtypes as can be seen in **Fig 3B**. For example, from the figure, strong positive and negative association of G25 status can be seen with Luminal A and basal-like subtypes respectively. Since G25 status has also shown strong association with ER and PR status its correlation with Luminal A (ER-positive, PR-positive and Her2 negative) and basal-like (triple negative) subtype is not surprising but highlights the versatility of gene group definitions.

Apart from PAM50 subtypes, we found the status of several gene groups associated with immune subtypes (C1, C2, C3 and C4) defined by Thorsson et al [27] as shown in **Fig 3B**. For example, from the figure strong association of Gene Group 15 (G15) can be seen with C2 (𝜌_r_ = 0.72, 𝑝 < 0.01) and C1 (𝜌_r_ = -0.48, 𝑝 < 0.01) and C3 (𝜌_r_ = -0.31, 𝑝 < 0.01). This association is expected as majority of G15 genes (*IFIT3*, *OAS3*, *IFI44L*, etc.) are interferon-regulated genes (IRGs) that play a role in the innate immune response and antiviral defense [28]. These results highlight the fact gene group statuses can be utilized as markers for immune activity as well as existing molecular subtyping of breast cancer patients.

#### 2.3.6 Association with mutations in cancer genes

We found the status of several gene groups associated with gene point mutation status (MUT) and copy number alteration status (CNA) as evident from **Fig 3B**. For example, from the figure, a strong negative correlation of G25 status with TP53 MUT status (𝜌_r_ = -0.59, 𝑝 < 0.01) and *MYC* CNA status (𝜌_r_ = -0.26, 𝑝 < 0.01) can be seen. Similarly, the status of several other gene groups can be seen as positively or negatively associated with MUT status (e.g., *CDH1, GATA3* and *PIK3A*) and CNA status (e.g., *ERBB2, PK2, HEY1, FGFR* and *F2F2*) of genes.

#### 2.3.7 Association of gene groups with pathologist-assigned histological phenotypes

We found gene groups status associated with routine clinical features such as histological types (invasive lobular and ductal carcinoma), histological grade (mitotic count, nuclear pleomorphism and epithelial tubule formation) [30] and the spatial fraction of tumor regions with tumor-infiltrating lymphocytes (TIL Regional Fraction) [31] as evident from **Fig 3B**. For example, from the figure, a positive correlation between G3 status and TIL Regional Fraction can be seen. Similarly, the status of G25 can be seen negatively associated with mitosis, necrosis, nuclear pleomorphism, inflammation and tumor grade, whereas positively associated with invasive lobular carcinoma. Association of G3 binary status with TIL Regional Fraction is expected as its status is defined by the expression level of several immune-related genes (e.g., *IL2, CD27, CCL5, PD-1* and *PD-L2*) [27], [32]. Similarly, G25 status negative association with mitotic count is not surprising as previous studies have found that over-expression of *MYC* (G25 = 1 when *MYC* is over-expressed) impairs mitotic spindle formation [33]. This analysis shows that gene group status can be associated with pathologist-assigned histological phenotypes.

### 2.4 Prediction of Gene Groups from histological imaging

To explore the association between phenotypic information contained in the WSI and the expression status of a set of genes in a certain gene group we have developed a novel deep learning based multi-task graph neural network pipeline (𝑆𝑙𝑖𝑑𝑒𝐺𝑟𝑎𝑝ℎ^oo^) that takes a WSI as input and predicts the status of 200 gene groups simultaneously. The workflow of the proposed approach is shown in **Fig 1B**. It builds on our previous work that can model a WSI as a graph to capture histological context but has been significantly expanded and improved [34].

#### 2.4.1 Quantitative results of prediction of individual gene group statuses

Our predictive analysis shows that the binary status of a significant number of gene groups can be predicted from histology images with high area under the receiver operating characteristic curve (AUROC). **Fig 4A** shows model performance in terms of mean AUROC. The binary status of many gene groups can be predicted with an AUROC of above 0.60. Additionally, the status of around 29 gene groups is predicted with a high AUROC of above 0.80. For the top 20 best-predicted gene groups we show the AUROC distribution across 1,000 bootstrap runs in **Fig 4B**. From the figure, G0, G100 and G25 status can be predicted with an AUC-ROC of above 0.87 with a narrow confidence interval.

**Figure 4.**
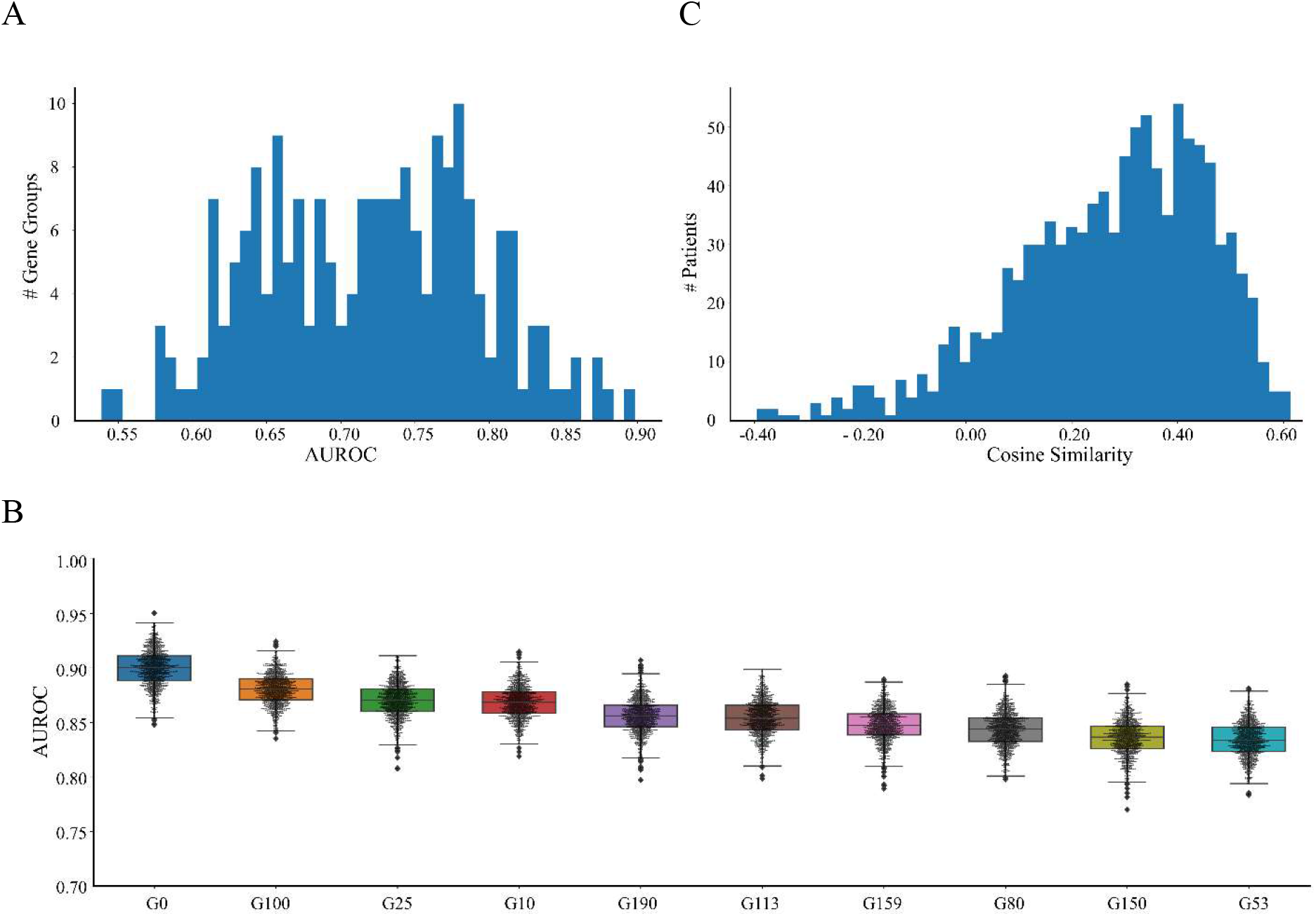
Quantitative result. A) Histogram displaying the AUROC at which the binary status of gene groups are predicted from WSIs. B) Box plot showing AUROC distribution of top-10 best predicted gene groups across-1,000 bootstrap runs. C) Histogram of patient-wise cosine similarity between true and predicted gene expression state.

To analyze the degree to which the complete gene expression profile of a patient can be predicted from imaging alone, **Fig 4C** displays a histogram of patient-wise cosine similarity between histology image-based inferred gene expression state and true gene expression state. From the plot, the similarity score shows a moderate alignment between the true and predicted gene expression states of each patient (average cosine similarity across all patients of 0.27). Of particular interest are patients whose alignment score is either very high or very low. Some example WSI thumbnails of patients whose expression state is best or poorly predicted from histological imaging are shown in **STable 1**. These results point to the fact that although the status of certain groups can be predicted with high accuracy, it is not possible to fully characterize the overall gene expression state of most patients from histological imaging alone. This result is expected due to both technical and underlying biological reasons. For example, histological imaging and gene expression analysis are carried out on different tissue sections and the latter uses “bulk” tissue. Furthermore, not all gene expression changes will have a phenotypic effect that can be observed in a WSI which in turn allows predictive modelling as illustrated in **SFig 7**. This shows that both whole slide imaging and gene expression analysis carry complementary value in understanding disease mechanisms.

#### 2.4.2 Spatial Profiling and histological phenotypes of Gene Groups

The proposed graph neural network can map WSI-level predictions of a gene group to spatially localized regions or nodes in the input image. This enables the profiling of local histological patterns linked to gene groups based on their node-level predictions. **Fig 5** shows the spatial profiling of gene groups (G3 and G25 as examples) by visualizing node-level prediction scores from 𝑆𝑙𝑖𝑑𝑒𝐺𝑟𝑎𝑝ℎ^oo^. For both gene groups, an example WSI with its corresponding heatmap highlighting node level prediction score is shown against binary status 0 and 1. The heatmap highlights the spatially resolved contribution of different regions of the WSI towards the expression status of a certain gene group being 0 or 1. More specifically, regions highlighted in redder color are indicative of an association with status = 1, whereas regions highlighted in bluish color are indicative of an association with status = 0 of a particular gene group. It is interesting to note that a given gene group exhibits significant variation in prediction score across different regions of the image, which can be linked to the spatial diversity of localized gene expression patterns throughout the tissue. The localized predictions for other gene groups can be viewed in the online portal (<u>HiGGsXplore</u>).

**Figure 5:**
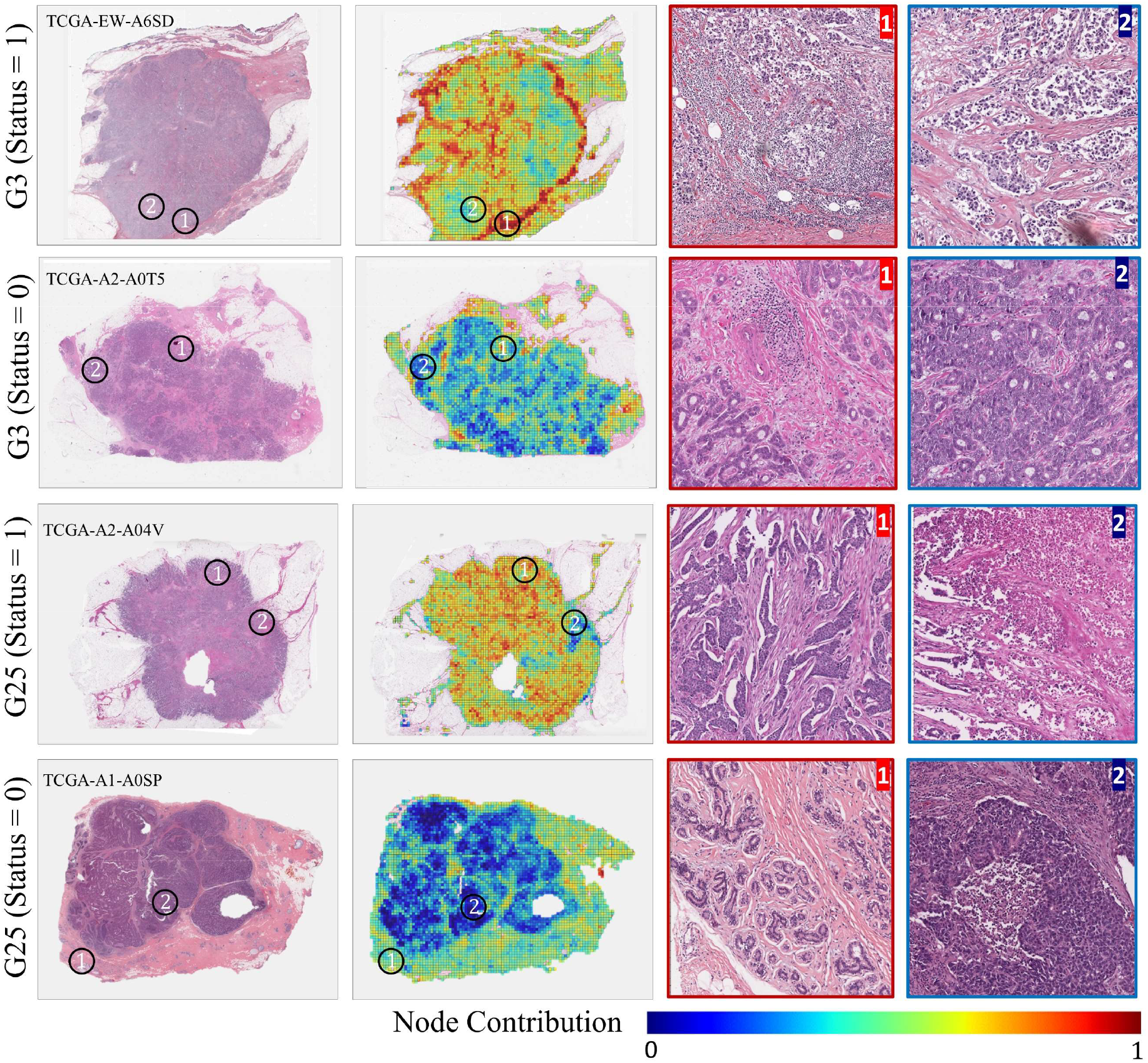
Spatial profiling of gene groups status. Spatial profiling of gene group 3 (G3) and 25 (G25) is displayed through example WSIs and heatmaps. The heatmaps use pseudo colors (bluish to red) to highlight the spatially resolved contribution of patches to the predicted expression state, with bluish and redder color indicating highly contributing status = 0 and status = 1 regions, respectively. From WSIs we extracted magnified version of highly contributing status = 0 and status = 1 regions (ROIs) outlined by red and blue color, respectively. The black circles highlight regions of WSIs from which ROIs were extracted. For an interactive visualization, please see: <u>tiademos.dcs.warwick.ac.uk/bokeh_app?demo=HiGGsXplore</u>

Using node-level prediction score as a guide, we extracted some regions of interest (ROIs) associated with G3 and G25 status = 0 and 1 from their corresponding WSI as shown in **Fig 5**. ROIs representative of G3 = 1 have a relatively high proportion of inflammatory cells compared to G3 = 0 ROIs where tumor cells appear more pleomorphic. Additionally, for the patient with G3 (status = 1), the invasive margin of the tumor, which has a higher density of inflammatory cells, is shown to be correlated with G3 status = 1. Given that G3 status is associated with TIL regional fraction (see Fig 3) and immune response related processes and pathways (see Fig 2C, SFig 2 and SFig 4), therefore tumor-infiltrating lymphocytes (TILs) is the likely histological phenotype associated with G3 (status = 1). This also explains the higher survival probability of G3 (status = 1) patients as several studies have found TILs associated with good prognosis [35]. Regarding G25, tubule formation, and normal lobule can be seen in ROIs representative of G25 (status = 1), whereas, in ROIs indicative of G25 (status = 0) the obvious feature is necrosis, and more pleomorphic tumor cells. For the patient with G25 (status = 1), regions of the WSI with tubule formation are highlighted as evident from the ROI. However, for patient with G25 (status = 0) tissue regions with normal lobule received higher score since there was no tissue area with tubule formation. The highlighted spatially resolved histological patterns are concordant with their corresponding enriched cancer hallmark processes (Estrogen response, Immune response and p53 signalling) and biological pathways (see Fig 2C, SFig 2 and SFig 4).

This analysis shows that the proposed deep learning pipeline has identified relevant spatially resolved histological patterns associated with different gene groups (TILs in the case of G3 and tubule formation in the case of G25) in an automated manner as evident from the heatmaps. It is noteworthy, that in cases where no tubule formation is present in the WSI (see G25 = 0 ROIs), it has highlighted normal lobule which is quite remarkable.

#### 2.4.3 Mining differential histological patterns associated with each gene group

To explore the association between visual patterns contained in WSIs and gene groups status we identified 25 exemplar patches for each status (0 and 1) of a certain gene group. For these patches, we also computed the cellular composition (counts of neoplastic, inflammatory, connective, and epithelial cells), overall cellularity and mitotic counts. **Fig 6A** shows 10 out of 25 representative patches for each of G3 and G25 status = 0 and status = 1. The main difference between G3 = 0 and 1 patches, as seen in the figure, is the presence of lymphoid infiltrate and tumor cellularity. More specifically, G3 = 1 patches have more inflammatory cells and fewer neoplastic cells, whereas the opposite is true for G3 = 0 patches. This differential histological pattern across all patients is concordant with the spatially resolved visual pattern we see in G3 = 0 and 1 ROIs (see Fig 5) and can be used as a histological motif. Additionally, G3 = 0 patches have relatively higher number of mitotic counts compared to G3 = 1. Regarding G25, the striking difference between G25 = 0 and G25 = 1 patches is the presence of tubule formation (row 2 patch 2 and 3, row 2 image 2 and 3) in the tumor area. As G25 status correlates positively with ER and PR status (see Fig 3B) and previous study has also found ER and PR positive cancers enriched in tubule formation [36], therefore, tubule formation could be the histological phenotype associated with G25 = 1. In contrast, G25 = 0 patches have more pleomorphic sheets of cells and areas of necrosis (row 1 image 1 and 3, row 2 image 1 and 2). This pattern agrees with the histopathological phenotypes we observed in **Fig 3B** and **Fig 5**. Finally, G25 = 1 patches show higher mitotic and inflammatory cell counts compared to G25 = 0 patches. Though we are not using any histopathological annotations in training, the predictive model has identified relevant morphometric patterns in an automated manner.

**Figure 6:**
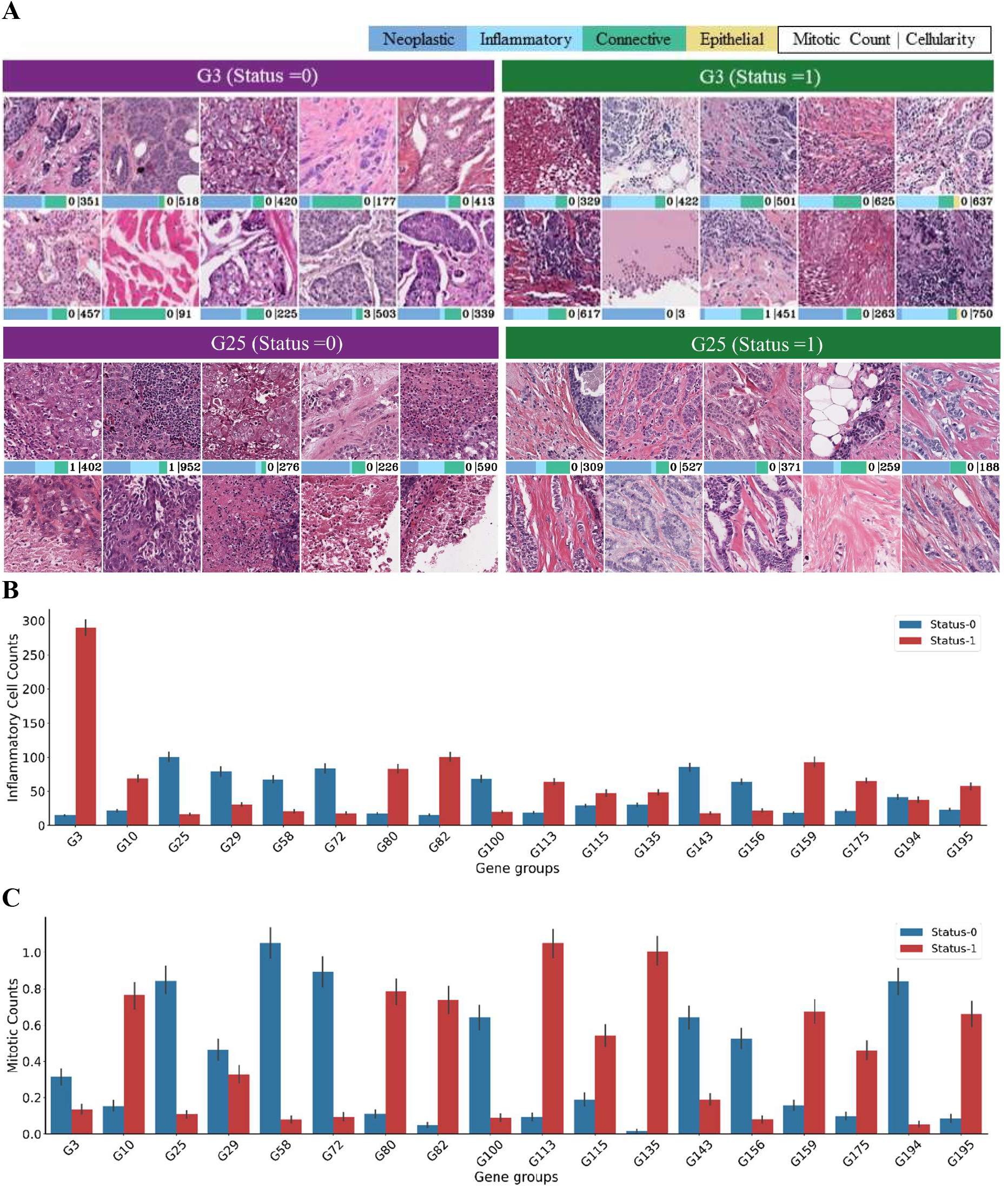
Histlogical patterns associated with gene groups. A) Representative patches of G25 and G3 status 1 and 0 are shown. The bar below the patches shows patch level cellular composition, mitotic counts and cellularity. B) Gene groups status (0 and 1) association with patch-level Inflammatory cell counts. C) Gene groups status (0 and 1) association with patch-level mitotic cell counts.

Apart from G25 and G3, we found patch-level inflammatory cell counts and mitotic counts statistically significantly associated (Wilcoxon test 𝑝 < 0.01) with the binary status of several other gene groups as shown in **Fig 6B** and **Fig 6C**.

### 2.5 Image-based predicted gene group statuses provide latent space for down-stream predictive modeling

Gene expression groups allow us to capture the gene expression profile of a given patient in terms of 200 gene status variables and their prediction through a machine learning model allows us to map histological patterns to these gene groups. However, the predicted statuses of gene groups can also be used as a compressed latent space representation for predictive modelling of other histologically important clinical variables. **Fig** (**7A-F**) show the predictability of clinical variables based on the predicted gene group statuses as latent variables using a simple linear classifier. PAM50 subtypes such as Basal, Luminal A, Luminal B and Her2 can be predicted from these latent variables with a mean AUROC of 0.90, 0.82, 0.78 and 0.75 respectively. Similarly, the latent representation can also predict the status of ER, PR and Her2 with a mean AUROC of 0.88, 0.79 and 0.61 respectively. Apart from this, we found the latent variables predictive of several signalling pathways alteration status, immune subtype, and also genes MUT status and CNA status. For example, TP53 pathway alteration status can be predicted with a mean AUROC of 0.75 from these latent variables [37]. The latent variables can also predict MUT status (14 genes) and CNA status (12 genes) with an AUROC of above 0.60 as evident from **Fig 7E** and **Fig 7F**. For example, TP53 point MUT status and *ERRB2* CNA status can be predicted with an AUROC of 0.81 and 0.79 respectively, which are higher that baseline results of 0.79 for *TP53* MUT status [38] and 0.62 for *ERBB2* MUT status [39]. **Fig 7G** shows some example heatmaps demonstrating spatial profiling of these clinical variables. From figures, ER and PR status have similar highlighted regions, while basal subtypes (ER, PR and Her2 negative) have opposite regions. The heatmaps also show the spatial profiling of Luminal B subtype, and TP53 MUT and pathway alteration status. This clearly illustrates the value of the proposed gene groups for downstream predictive modelling.

**Figure 7:**
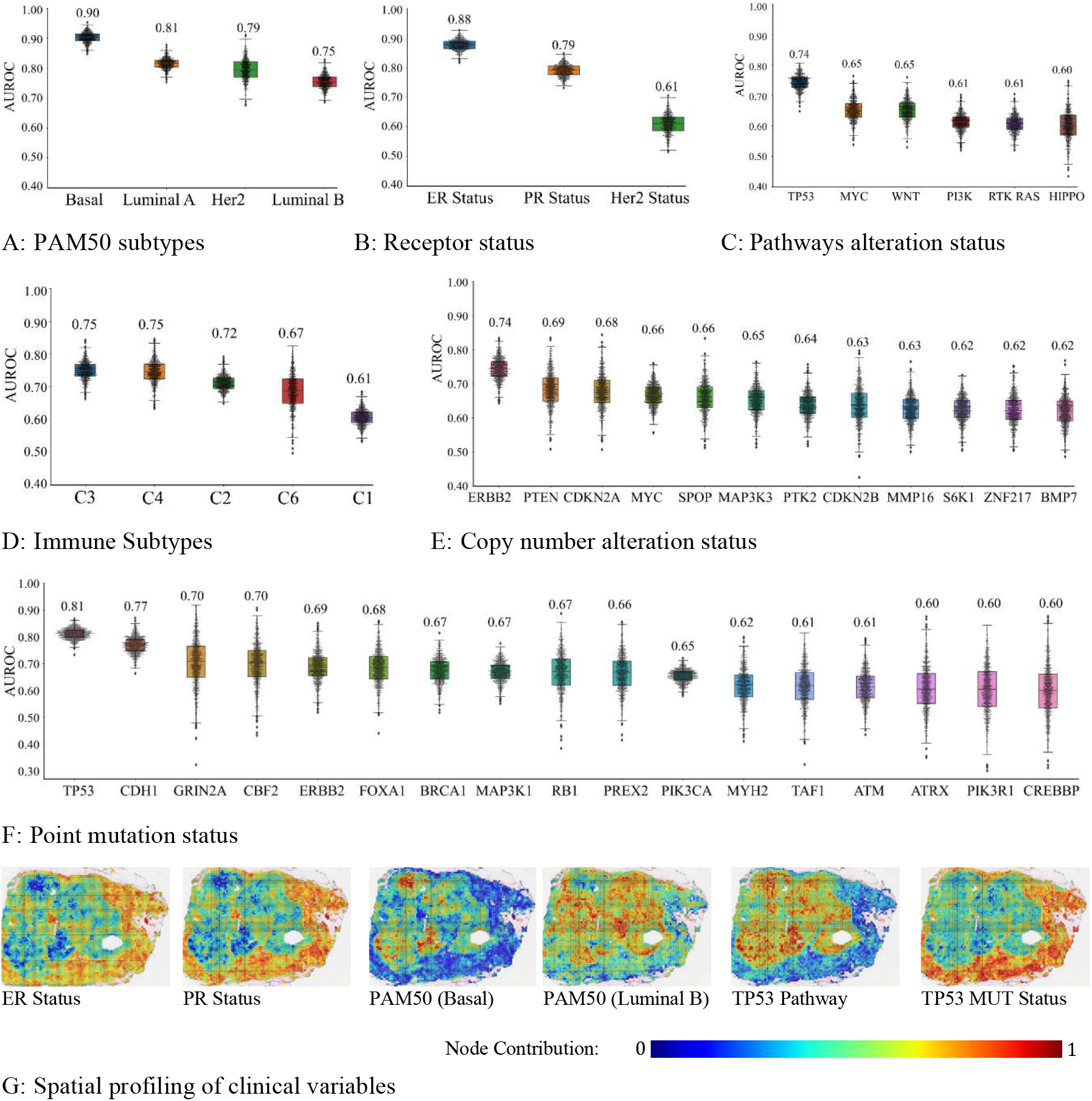
Implication of Image-based predicted gene group statuses for downstream predictive modeling. Prediction of (A) receptor status, (B) PAM50 molecular subtypes, (C) Immune subtypes, (D) pathways alteration status, (E) driver genes copy number alteration status and (F) point mutation status from image-based predicted gene groups status. Each box in the figure shows the AUROC distribution at which a clinical variable is predicted from image-based predicted gene group status across 1, 000 bootstrap runs. The scatter plot on top of box plot shows the AUROC values across different bootstrap runs while the numeric value above each box shows the mean AUROC value. G) Spatial profiling of some routine clinical variables is shown using example heatmaps. The heatmaps use pseudo colors (bluish to red) to highlight the spatially resolved contribution of patches to status = 0 and 1 of a certain clinical variable, with bluish color indicating highly contributing status = 0 regions and red color indicating highly contributing status = 1 regions.

### 2.6 Clinical and Therapeutic significance of best-predicted gene groups

We found that gene groups predicted with high accuracy (AUROC ≥ 0.75) from imaging are significantly associated with disease specific survival (DSS), biological pathways and hallmark processes. All 25 gene groups associated with DSS are predicted with high accuracy from imaging. Besides this, some interesting biological pathways (see **Fig 8**) and cancer hallmark processes (see **SFig 8**) can also be inferred from images based predicted gene groups which can guide histology image-based therapeutic decisions by selecting drugs that target a certain biological pathway (e.g. PI3K-Akt) [40].

**Figure 8:**
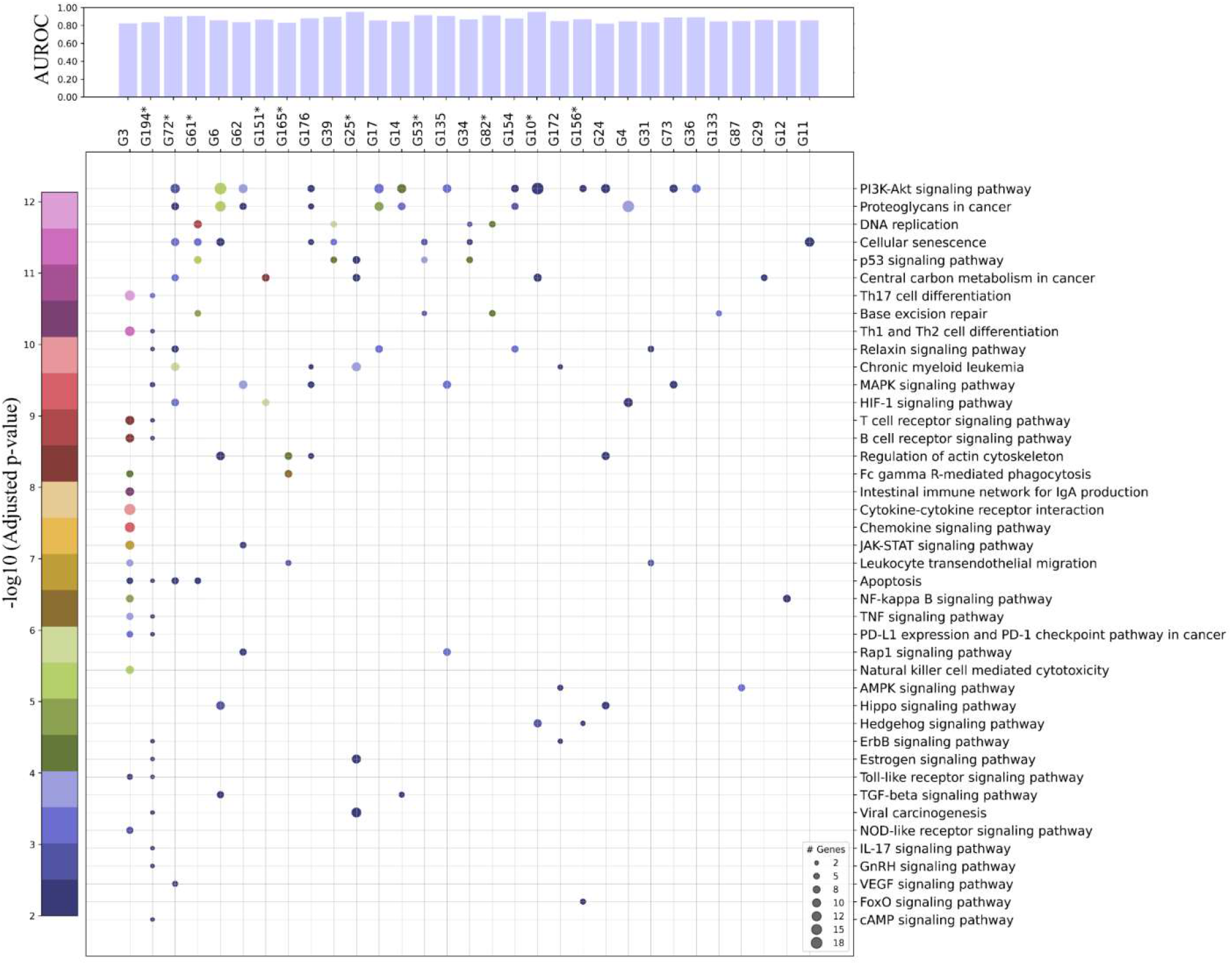
Clinical and Therapeutic significance of best predicted gene group. The scatter plot shows association of gene groups with biological pathways with gene group shown along x-axis (one per column) and corresponding enriched pathways on y-axis (one per row). The size of scatter shows the number of genes from a particular gene group that has shown statistically significant association (FDR adjusted p-value < 0.01) with a certain biological pathway. In the plot the p-value is represented by the color of scatter dots. The top bar plot shows the prediction accuracy (AUROC) at which the status of these gene groups is predicted from histology images. Gene groups that show statistically significant association with disease specific survival are annotated with a * next to the gene group name.

## 3 Discussion

We performed histological and molecular characterization of breast cancer patients using a purely data-driven approach. Highlighting the limitations of previous methods that predict the expression level of individual genes from histology image, we have shown that significant co-dependencies of different genes across samples (see **Fig 2B**) compromises the ability of deep learning models to identify individual gene level genotype to phenotype mapping. To tackle this, we first grouped genes whose expression patterns are significantly dependent and covarying across samples and then proposed a multi-output graph-based deep learning pipeline (𝑆𝑙𝑖𝑑𝑒𝐺𝑟𝑎𝑝ℎ^oo^) that predicts both WSI-level and spatially resolved expression status of these gene groups in an end-to-end manner. Using the proposed computational pathology workflow, we demonstrated that the status of a significant number of gene groups can be predicted with high accuracy from imaging. This not only overcomes the limitations of existing image-based gene expression prediction models but provides opportunities to gain biological insights from imaging directly. Finally, we showed that histopathological patterns associated with several gene groups in terms of cellular composition, mitotic counts and exemplar patches can be identified using the proposed computational pathology pipeline.

A potential advantage of the employed gene grouping approach is the interpretability of gene groups. The method allows a compact representation of a patient’s gene expression state (200 binary latent variables) without losing interpretability, which is crucial in this context as it provides insight into biological processes and underlying protein-protein and also drug-protein interactions that can motivate new therapies. Through GSEA, we found genes from several gene groups associated with cancer hallmark processes (e.g. EMT, inflammatory response, estrogen early and late response, mTORC1 signalling, Myc targets, p53 signalling, KRAS up and down signaling) and biological pathways (e.g. Inflammatory response, PD-L1 expression and PD-1 checkpoint, cancer immunotherapy by PD-blockade and EGF/EGFR signalling). Additionally, we have shown that genes in a certain gene group are enriched for protein-protein interaction that can be used for the identification of drugs that modulate the activity of a target protein of interest which will subsequently lead to precise diagnosis of patient tumor.

Another important observation regarding gene grouping is that, though the gene groups are defined in a completely data-driven manner without any intelligent selection still they carry significant clinical meaning in terms of association with survival (OS, DSS and PFS), routine clinical biomarkers (ER, PR and Her2 status), driver genes mutation statues, and previously defined PAM50 and Immune subtypes. Apart from this, we found the binary status of several gene groups associated with histopathological annotations which enable direct genotypic to phenotype mapping. Additionally, this genotype to phenotype link can further be validated using GSEA and specialized IHC staining. These results not only validate the clinicopathological significance of these gene groups but also provide a broader picture of an individual tumor by illuminating the interplay between patient gene expression state and several other clinical variables of interest.

A striking feature of the proposed approach for mapping patient gene expression status with morphometric patterns contained in the WSIs is its reliability and explainability. Localized histological patterns identified by 𝑆𝑙𝑖𝑑𝑒𝐺𝑟𝑎𝑝ℎ^oo^can be explained in terms of enriched hallmark process, biological pathway and underlying protein-protein interaction, and also through specialized IHC staining and genome sequencing. For example, we found genes from G3 enriched for several immune-related biological processes and pathways including PD-L1 expression and PD-1 checkpoint pathway which in histology images we found associated with a high proportion of TIL. Thought the observation is interesting but still further validation is needed using IHC data. After validation, this will allow the selection of patients for immunotherapy based on routine histology images. Regarding G25 we found tubule formation in majority of G25 = 1 representative patches, which was consistent with IHC ER and PR status and also the associated cancer hallmark process (Estrogen signalling). In contrast, G25 = 0 patches have more pleomorphic sheets of cells several with area of necrosis, which is again concordant with their association with pathologist-assigned phenotypes (necrosis and nuclear pleomorphism), TP53 MUT status and p53 signalling pathway. This show that the proposed deep learning pipeline has identified relevant spatially resolved histological patterns associated with the status of gene groups in an automated manner.

Image-based prediction of gene expression state will open doors of gaining biological insights from imaging directly and is expected to be advantageous in both cancer research and clinical setup. In cancer research, the proposed approach can be used for studying the interplay between gene expression and histopathological phenotypes. Additionally, it can also be used by pharmaceutical industries in their drug discovery pipeline when they study the response of lead compounds in early-phase trials. In clinical setup, it will allow cost-effective precision diagnostic from imaging data alone. The proposed computational pathology pipeline not only predicts patient gene expression but also provides a detailed insight in terms of patient survival (OS, DFS and PFS), possible up or downregulated biological processes and their underlying protein-protein interaction, possibly mutated or copy-altered genes, and information about ER, PR and HER2 status, PAM50 and immune subtypes. These types of analysis will provide a more detailed insight into an individual tumor in a cost-effective way. It is important to highlight here, that though we managed to predict the expression status of several gene groups with high accuracy and we extensively validated the results, further extensive validation on a large multi-centric dataset is needed before entering into clinical practice.

## Acknowledgments

M.D. would like to acknowledge the PhD studentship support from GlaxoSmithKline. F.M and N.R supported by the PathLAKE digital pathology consortium which is funded from the Data to Early Diagnosis and Precision Medicine strand of the government’s Industrial Strategy Challenge Fund, managed and delivered by UK Research and Innovation (UKRI). F.M and M.E also acknowledge funding support from EPSRC EP/W02909X/1.

## Author Contributions

Conception: FUAAM, NR, KB and MD; Experiment Design: MD, FUAAM, NR, KB, ABH; Bioinformatics analysis: FUAAM, ABH, MD; Pathologist review: LJ; Clinical Review: LJ and LY; Coding and data analysis: MD; Visualization and portal development: ME and MD; Mitotic data analysis: MJ and MD; Writeup: MD and FM with input and review from all authors; Funding acquisition: NR, FUAAM and KB.

## 4 STAR Methods

### 4.1 Dataset

#### 4.1.1 Acquisition and preprocessing of RNA-Seq data

We collected RSEM (RNA-Seq by Expectation and Maximization) normalized RNA-Seq data of 1084 TCGA breast cancer patients from cBioportal [41], [42]. The gene expression data was obtained using log2 normalized z-score values of the expression of 5,596 genes having high variance in expression across patient samples along with known oncogenes.

##### 4.1.1.1 Acquisition of whole slide images and survival data

We collected 1,133 Whole Slide Images (WSIs) of Formalin-Fixed Paraffin-Embedded (FFPE) Hematoxylin and Eosin (H&E) stained tissue section of 1084 patients having breast cancer from the Cancer Genome Atlas (TCGA) [43], [44]. For patients with multiple slides, we selected the one with best visual quality. Additionally for robust analysis, we ignored WSIs with missing baseline resolution information. After slide filtering, we used 1,050 WSIs each belonging to an individual patient to avoid any overlap between training and testing over the same patient. For these patients, we used the survival data from the TCGA standardized clinical dataset called Pan-Cancer Clinical Data Resource (TCGA-CDR) [45] and other clinical data from cBioportal. For these patients we obtained annotation of 11 histopathologic features scored by pathologist from the data released by Thennavan et.al [30].

### 4.2 Data Driven Discovery of Gene Groups with CorEx

To model associations between expression profile of different genes we used Total Correlation Explanation (CorEx) on the gene expression matrix 𝑀 of size 𝑚 × 𝑛 where 𝑚 and 𝑛 are the number of patient samples, and genes, respectively [46]. As the expression of different genes is significantly inter-dependent and correlated, CorEx allows us to represent the gene expression state of a patient in terms of a small number of binary variables or gene groups that can capture information contained in the expression of all genes of a given patient with minimal loss. For a detailed mathematical formulation underlying CorEx, the interested reader is referred to the CorEx paper [46]. Given 𝑀_m×n_ as input, the output of CorEx is a matrix 𝐺_m×d_ with each column of 𝐺 corresponds to a binary latent factor 𝐺_k_ (𝑘 = 1 … 𝑑 with 𝑑 ≪ 𝑛) so that the mutual information between the expression level of genes is minimized after conditioning on 𝐺_l_, …, 𝐺_d_. In other words, the latent factors identified by CorEx can *“explain away”* the association between expression of various genes. Akin to “loadings” in principal component analysis (PCA), the definition of each binary latent factor 𝐺_k_ is based on mutual information between the expression score of a certain gene and the binary status of 𝐺_k_ across patient samples. This allows us to model each of the latent factors as a ranked (by mutual information) collection or group of genes. However, unlike PCA (or other linear or kernelized dimensionality reduction techniques based on covariance), CorEx can capture non-linear statistical relationships and dependencies between input variables (genes) directly due to its use of mutual information (see comparative analysis in [46]). Furthermore, CorEx produces binary latent factors which can be easier to interpret as the status of a certain gene group for a given patient will either be 0 or 1. We run the algorithm for 100 iterations on the z-score expression of TCGA-BRCA patients for discovering 200 binary latent factors. The number of latent factors were decided based on the TC distribution shown in **SFig 9**. The distribution demonstrates that the overall TC (sum of TCs of all latent factor) plateaus and approaches zero after selecting 200 latent factors. Therefore, we selected 200 latent factors. The binary statuses of these 200 latent variables define the expression state of a patient, where the binary value of each latent variable is defined by the group of genes whose gene expression patterns are substantially co-dependent across samples as shown in **Fig 2B**.

#### 4.2.1 Analysis of Biological and Therapeutic Significance of gene groups

Hallmark processes and KEGG pathways enrichment for genes in different gene groups were obtained using Enrichr [47]. In line with previous work [20], we selected a maximum of top 400 genes from each gene group whose mutual information is greater than 0.002. We passed the gene set to Enrichr which returns the enriched terms across a selected library (in our case KEGG pathway and MSigDB hallmarks) coupled with their statistical significance (FDR-adjusted p-value using Benjamini-Hochberg methods). We used a cutoff value of 𝑝 < 0.01 on the adjusted p-value for statistical significance of an enriched term across the selected library. The protein-protein and drug-protein interactions are analyzed using STITCH [48].

### 4.3 WSI Analysis Pipeline with *SlideGraph*^∞^

#### 4.3.1 Preprocessing of whole slide images

We segment the tissue regions of WSIs using a tissue segmentation model and ignore regions with tissue artefacts (pen-marking, tissue folding, etc.). Each WSI is then tiled into patches of size 512 × 512 pixels at a spatial resolution of 0.50 microns-per-pixel (MPP). Patches capturing less than 40% of informative tissue area (pixels with intensity higher than 200) are discarded, and the remaining patches (both tumor and non-tumor) are used.

#### 4.3.2 WSI-graph Construction

A graph = (𝑉, 𝐸) is defined by a vertex set 𝑉, and an edge set 𝐸. The set 𝑉 = {𝒗_𝒊_|𝑖 = 1, … 𝑁} defines nodes in a graph (in our case is the set of patches in a WSI) while connectivity between nodes is defined by the edges 𝐸. Each node 𝒗_𝒊_ = (𝒈_𝒊_, 𝒉_𝒊_) captures the spatial location (𝒈_𝒊_), and feature representation (𝒉_𝒊_) of a patch in the WSI. We obtain the feature representation 𝒉_𝒊_ ∈ 𝓡^𝟏𝟎𝟐𝟒^of a patch 𝑥_i_ by extracting latent representation from ShuffleNet [49] pretrained on ImageNet [50]. The edge set 𝐸 is obtained by connecting nodes to the neighboring nodes (distance less than 4000 pixels) using Delaunay triangulation. If two nodes 𝑣_i_ and 𝑣_j_ are connected, then there will be an edge 𝑒_ij_∈ 𝐸.

### 4.4 Gene expression state prediction using Graph Neural Network

We pass the graph representation of a WSI through a Graph Neural Network (GNN) for predicting the node-level and WSI-level expression status of all gene groups simultaneously. In this work, we have developed a custom multi-output GNN that predicts the patch-level and WSI-level expression statuses of different gene groups in an end-to-end manner. Node level representation is passed through EdgeConv layers 𝐿 = {1,2,3}. Each EdgeConv layer [51] updates the representation of each node in the graph by aggregating the information from their neighboring node and generates embedding for successive layers. For a node in layer 𝑙 at index 𝑚 the output embedding of EdgeConv layer can mathematically be written as follows:

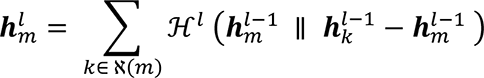

In the above equation 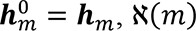 represents the neighboring nodes of 𝑚, and ℋ^l^ denote a neural network. EdgeConv operation is trying to combine information of a node 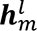 and neighboring nodes ℵ(𝑚). Since we are using three EdgeConv layers, each node is expected to capture information from the neighboring nodes that are less than 5-hops apart in the WSI-graph.

For spatial profiling for gene expression groups, the feature representation 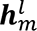 of a node 𝒗_m_ = (𝒈_𝒋_, 𝒉_𝒋_) ∈ 𝑉 is passed as input to a multilayer perceptron 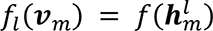 for generating node level prediction score which is then aggregated across all layers for getting patch level prediction score for all gene groups.

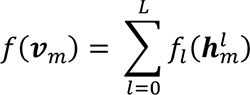

The WSI-level score for the expression status of all gene groups is obtained by pooling and aggregating node-level prediction scores as follows:

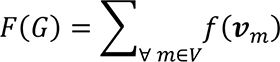

The trainable parameters of the EdgeConv layers and node-level classifiers are learned in an end-end manner using backpropagation. In a training batch of size 𝑁, the model predicted score for 𝑘 = {1 … 𝐾} binary latent factors are compared with their ground truth value using pairwise ranking loss [34], mathematically formulated as follows:

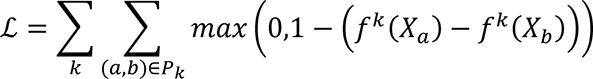

Here 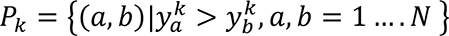 is the set of all pair of patients (a, b) where the expression status of patient 𝑎 is greater than patient 𝑏 for latent factor 𝑘. Minimization of the loss function ℒ(∵) will enforce the model to rank status = 1 patients higher than status = 0 for all latent factors.

### 4.5 Training and evaluation of *SlideGraph*^∞^

We trained and evaluated the performance of 𝑆𝑙𝑖𝑑𝑒𝐺𝑟𝑎𝑝ℎ^oo^ using 5-fold cross-validation, in which the dataset is subsampled into five 80/20 non-overlapping splits. The model is trained on 80% of the data and 20% data is held out for testing. From the training data we randomly select 10% of the data for parameter tuning and optimization. We train 𝑆𝑙𝑖𝑑𝑒𝐺𝑟𝑎𝑝ℎ^oo^ on the training set for 300 epochs using the Adam optimizer with an initial learning rate and weight decay of 0.001 and 0.0001, respectively. In each epoch, the training set is sampled into mini-batches of size 8, and the learnable parameters of 𝑆𝑙𝑖𝑑𝑒𝐺𝑟𝑎𝑝ℎ^oo^ are updated using adaptive momentum based optimizer. To avoid overfitting, we stop the model training early, if performance over the validation set does not improve for 20 consecutive epochs. During training, we maintain a queue of size 10 for tracking the best models based on their performance over the validation set. More specifically, we insert the model into the queue if the validation loss at epoch 𝑛 is less that the loss at epoch 𝑛 − 1. For test set inference, we ensemble the prediction score of all the models in the queue by averaging the prediction score and using that as the final prediction. For quantitative performance assessment, we report area under the receiver operating characteristic curve (AUROC) over the test set.

### 4.6 Spatial Profiling of Gene Groups and visualization

For a given WSI, the spatially resolved contribution of different tissue regions toward the expression status of a certain gene groups can visualized. We developed an online portal (http://tiademos.dcs.warwick.ac.uk/bokeh_app?demo=HiGGsXplore) which can assist user in spatially resolved cross-linking of genotype-phenotype mapping in terms of these gene groups. More specifically, the portal uses WSI couped with node level prediction of different gene group and then show the node level prediction in the form of an interactive heatmap. Additionally, the tool can also show different histological features when the user hover over a node in the graph.

### 4.7 Identification of Histological motifs

To uncover cellular and morphometric patterns associated with the expression status (0, or 1) of a particular gene group we divided patients into two groups (status = 0 and status = 1). For each group, we select 50 patients whose expression statuses are accurately predicted from their WSIs. From each of these WSIs, for patients with status = 1, we extract the highest scoring (based on node-level score) 1% patches, while for status = 0, we extract the lowest scoring patches and then cluster the patches within each group for getting representative patterns. Within each group (status = 0, and 1) we cluster the patches using 25-medoid clustering. After clustering, we get 25 visual patterns (histological motifs) representative of expression status = 0 and status =1 of a certain gene group.

### 4.8 Cellular composition estimations

We estimated the counts of neoplastic, inflammatory, connective, and normal epithelial cells present in a patch using our in-house cellular composition predictor ALBRT. ALBRT takes a patch of size 256 × 256 at a spatial resolution of 0.25 MPP and predicts the counts of the aforementioned types of cells present in it. We extracted patches of size 256 × 256 at 0.25 MPP using (x, y) of coordinates of 512 × 512 at 0.50 MPP. For each 512 × 512 patch, we obtained the cellular composition estimates by aggregating ALBRT-predicted cellular estimates of around 16 256 × 256 patches. The cellularity was computed by summing the counts of neoplastic, inflammatory, connective and epithelial cells present in a 512 × 512 patch.

### 4.9 Estimation of mitotic counts

Mitosis detection has been done using the state-of-the-art “mitosis detection: fast and slow” (MDFS) method [52]. MDFS is a two-stage method where mitotic candidates are first detected using a fully convolutional neural network and then refined by a deeper CNN classifier. Several techniques have been incorporated during the training of the MDFS to make it robust against domain shift problems seen in histology images and generalize better to unseen images. After detecting mitotic figures, we estimate the patch-level mitotic counts by counting all the detected mitoses in the patch.

### 4.10 Training and evaluation of Downstream predictors

We train separate multi-output perceptron for predicting the receptor status, PAM50 molecular subtypes, Immune subtypes, pathways alteration status, genes point mutation status and copy number alteration status using 𝑆𝑙𝑖𝑑𝑒𝐺𝑟𝑎𝑝ℎ^∞^ predicted gene groups status as features. The classifier for each downstream task is trained and evaluated using same loss function and training and validation protocol employed for 𝑆𝑙𝑖𝑑𝑒𝐺𝑟𝑎𝑝ℎ^oo^training and evaluation. After cross-validation, we get the downstream classifier prediction score for a particular clinical variable of interest for all patients. For performance we subsample 67% of the patients 1,000 times with replacement, and compute the AUROC between ground truth and model predicted score.

## Data and code availability

Whole slides images (WSIs) and corresponding genomic data and clinical data of all TCGA patients used in the study can be downloaded from NIH Genomic Data Common Portal at this link: https://portal.gdc.cancer.gov/. All genomic and histological analysis was performed in python. The deep learning model 𝑆𝑙𝑖𝑑𝑒𝐺𝑟𝑎𝑝ℎ^oo^ was developed using PyTorch Geometric library. Code and documentation of all python script used in the study can be found at: https://github.com/engrodawood/HiGGsXplore.

## Supplementary Materials

**SFig 1:**
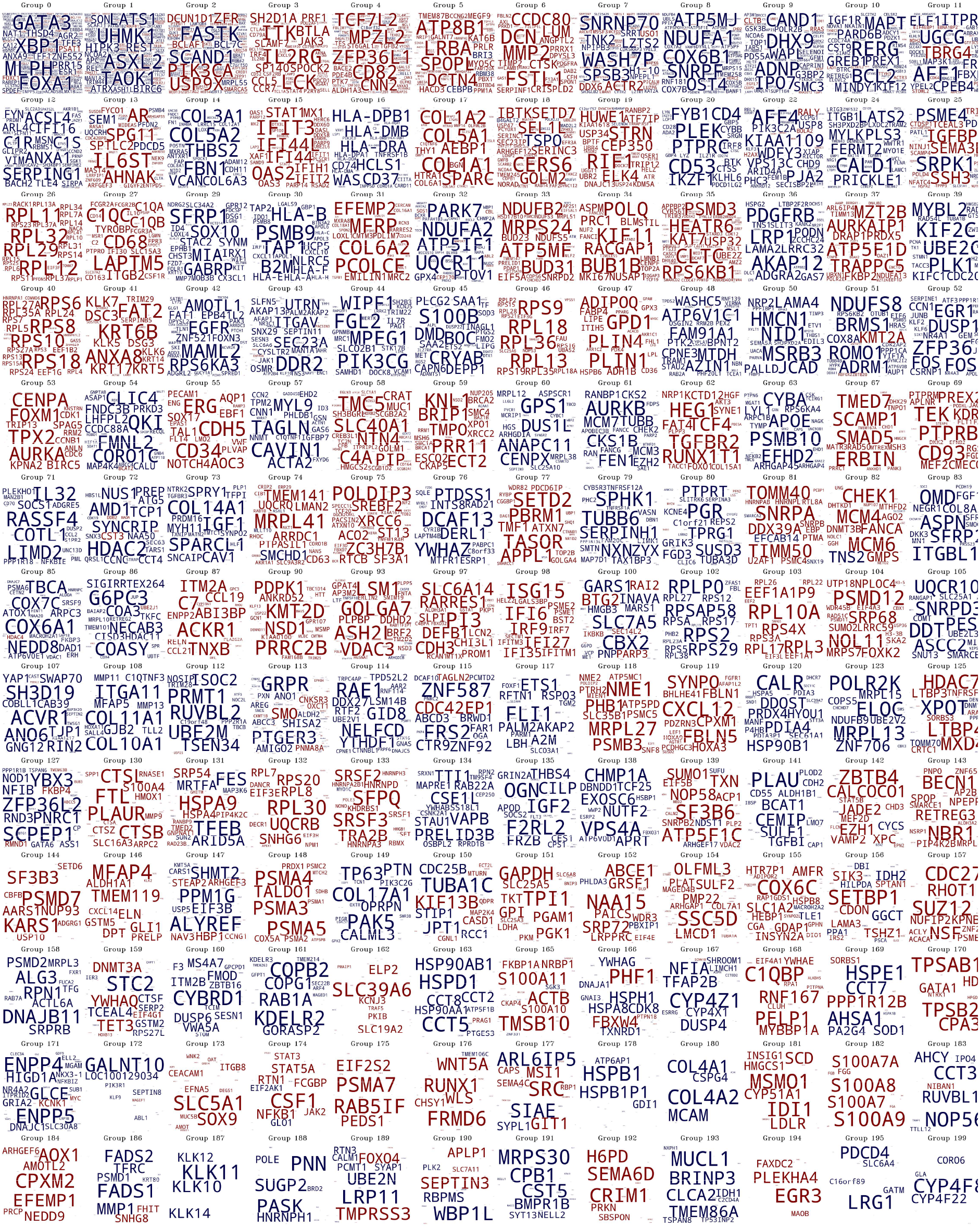
Word clouds demonstrating the gene composition of different gene groups. The color of the gene indicates whether its median expression across patients is high (red) or low (blue) when gene group status = 1. The font size of gene within a group is proportional to the amount of information that the gene status provides about a particular gene.

**SFig 2:**
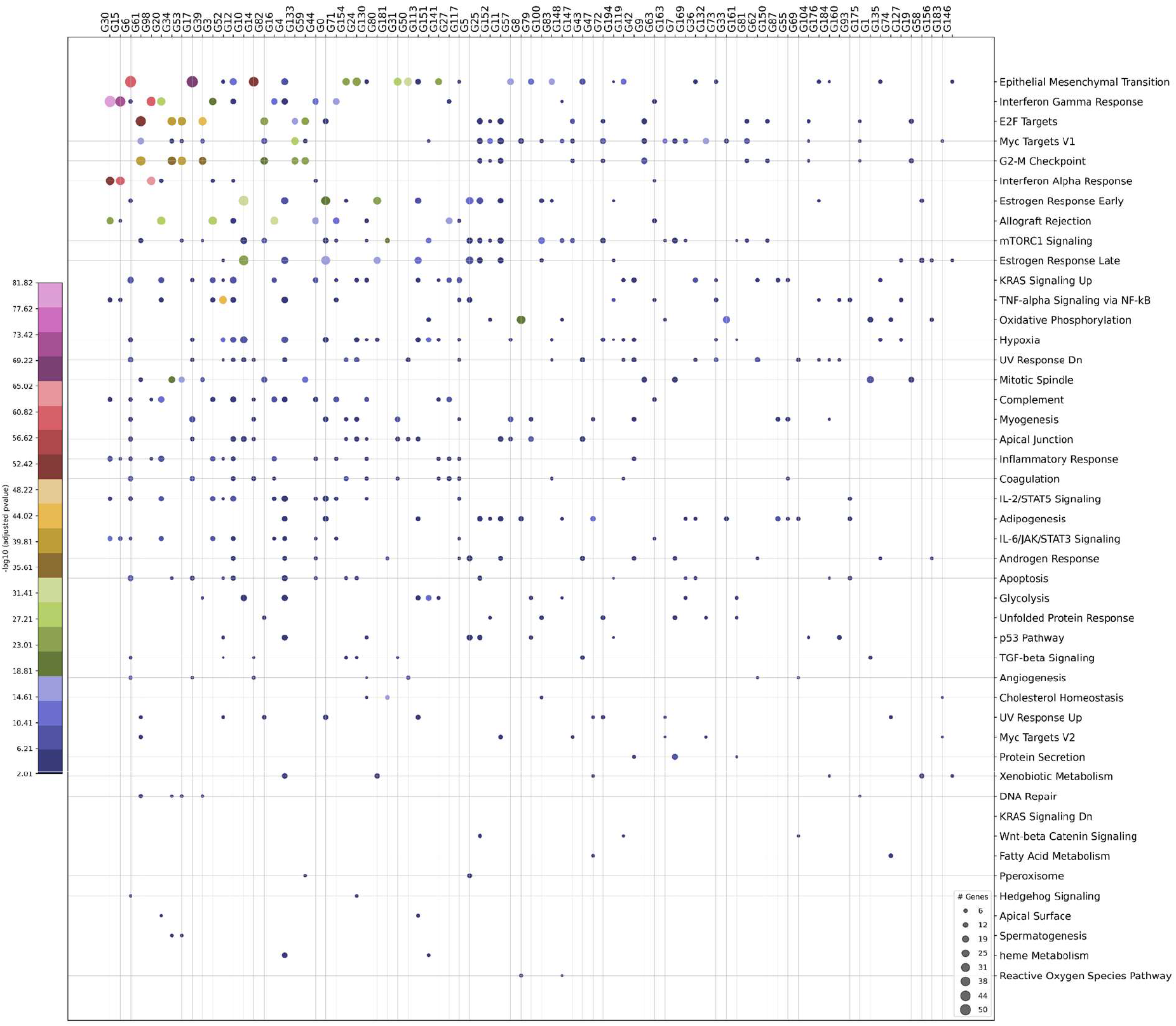
Enrichment of gene groups for cancer Hallmark processes is illustrated as 2D scatter plot with the gene group displayed along x-axis and the corresponding enriched biological pathways on y-axis. The size of the dot represents the number of genes from a specific gene group that has shown enrichment for a particular hallmark process while its color represents the statistical significance of association in terms of FDR-corrected p-value.

**SFig 3:**
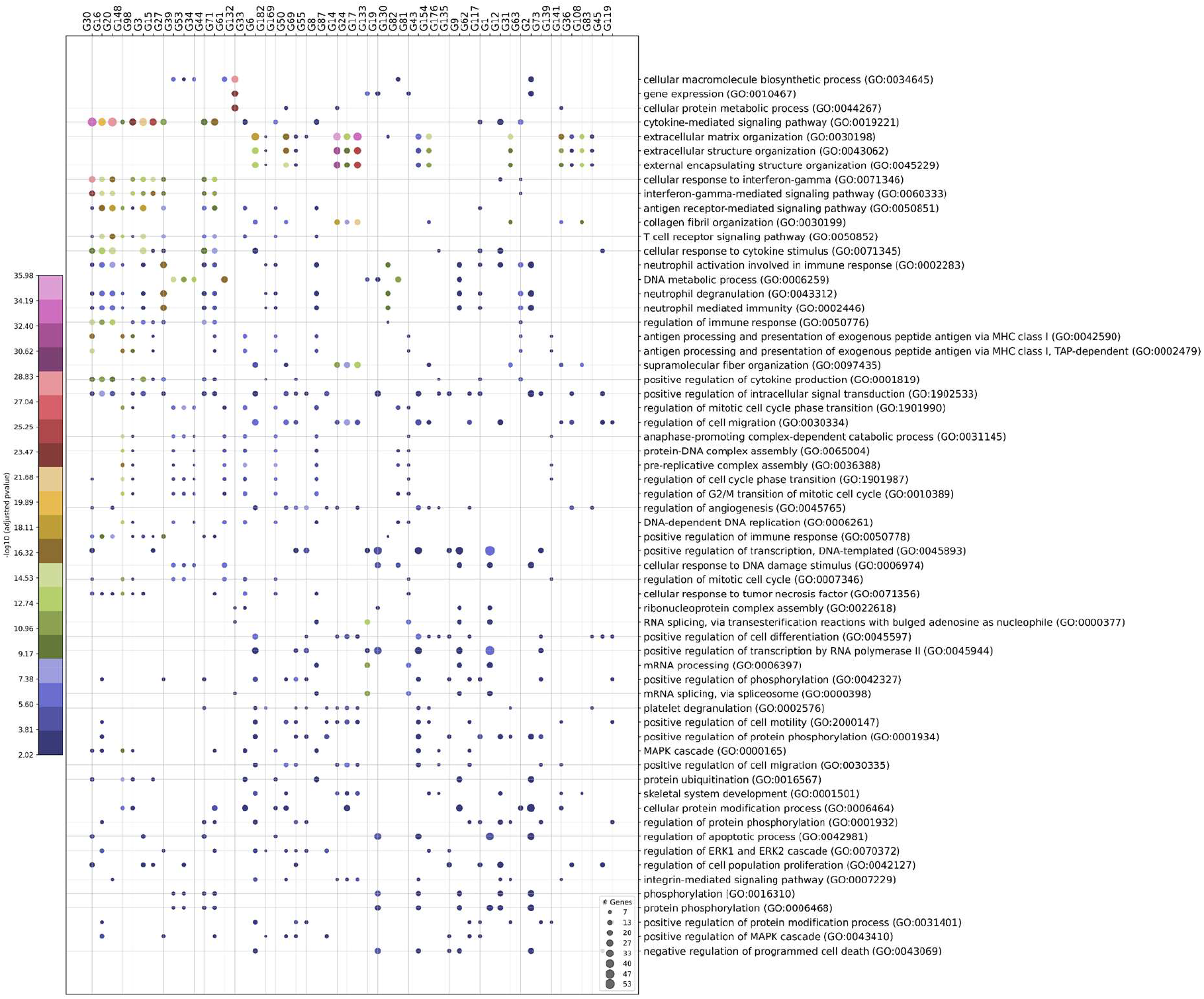
Enrichment of gene groups for GO (Gene ontology) biological processes is shown as 2D scatter plot with the gene groups displayed along x-axis and the corresponding enriched biological processes on y-axis. The size of scatter dot represents the number of genes from a specific gene group that has shown enrichment for a particular biological process while its color represents the statistical significance of the association in terms of FDR-corrected p-value.

**SFig 4:**
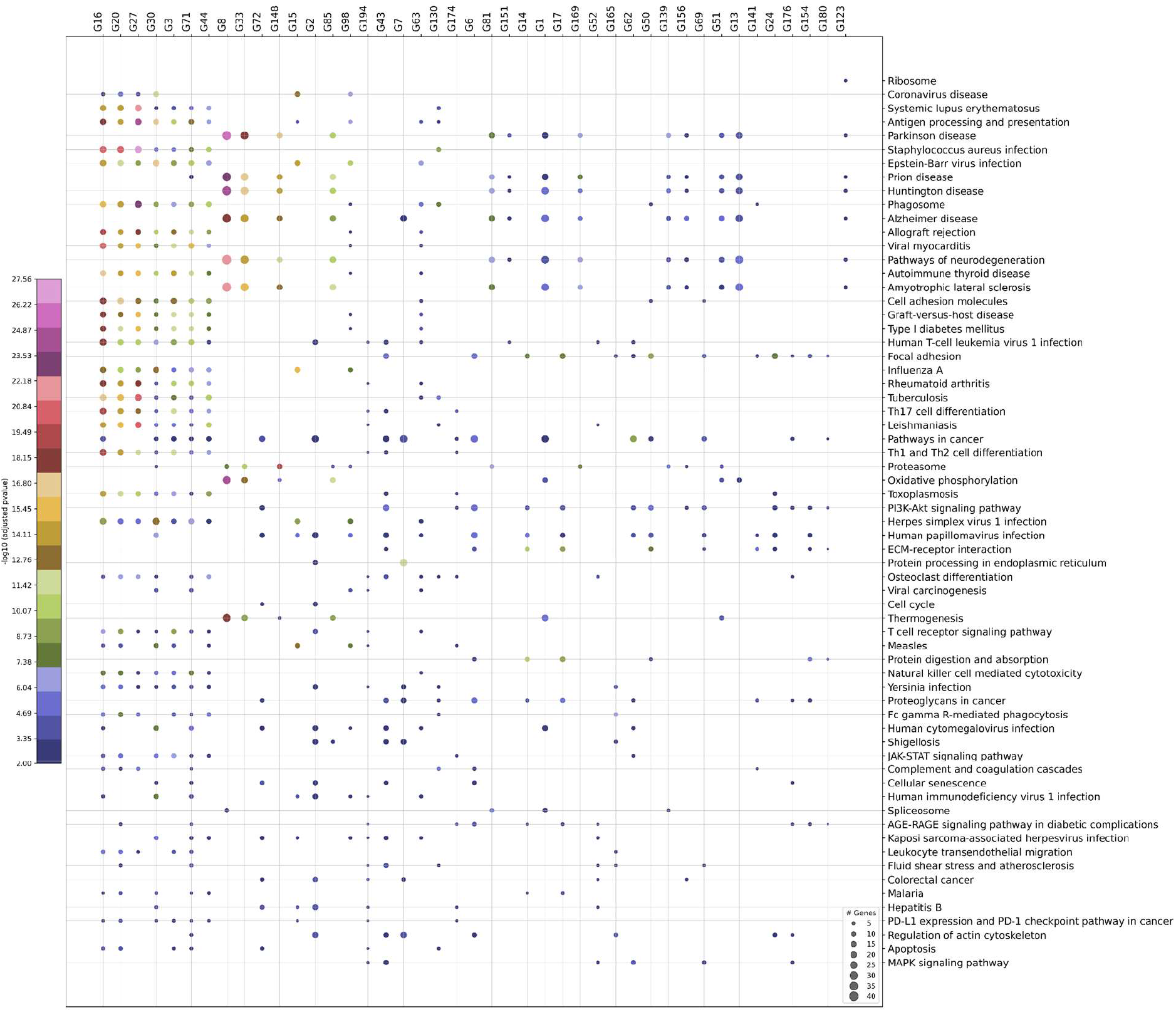
Enrichment of gene groups for KEGG Pathways is presented as 2D scatter plot with the gene group displayed along x-axis and the corresponding enriched biological pathways on y-axis. The size of the dot represents the number of genes from a specific gene group that has shown enrichment for a particular biological pathway while its color represents the statistical significance of association in terms of FDR-corrected p-value.

**SFig 5:**
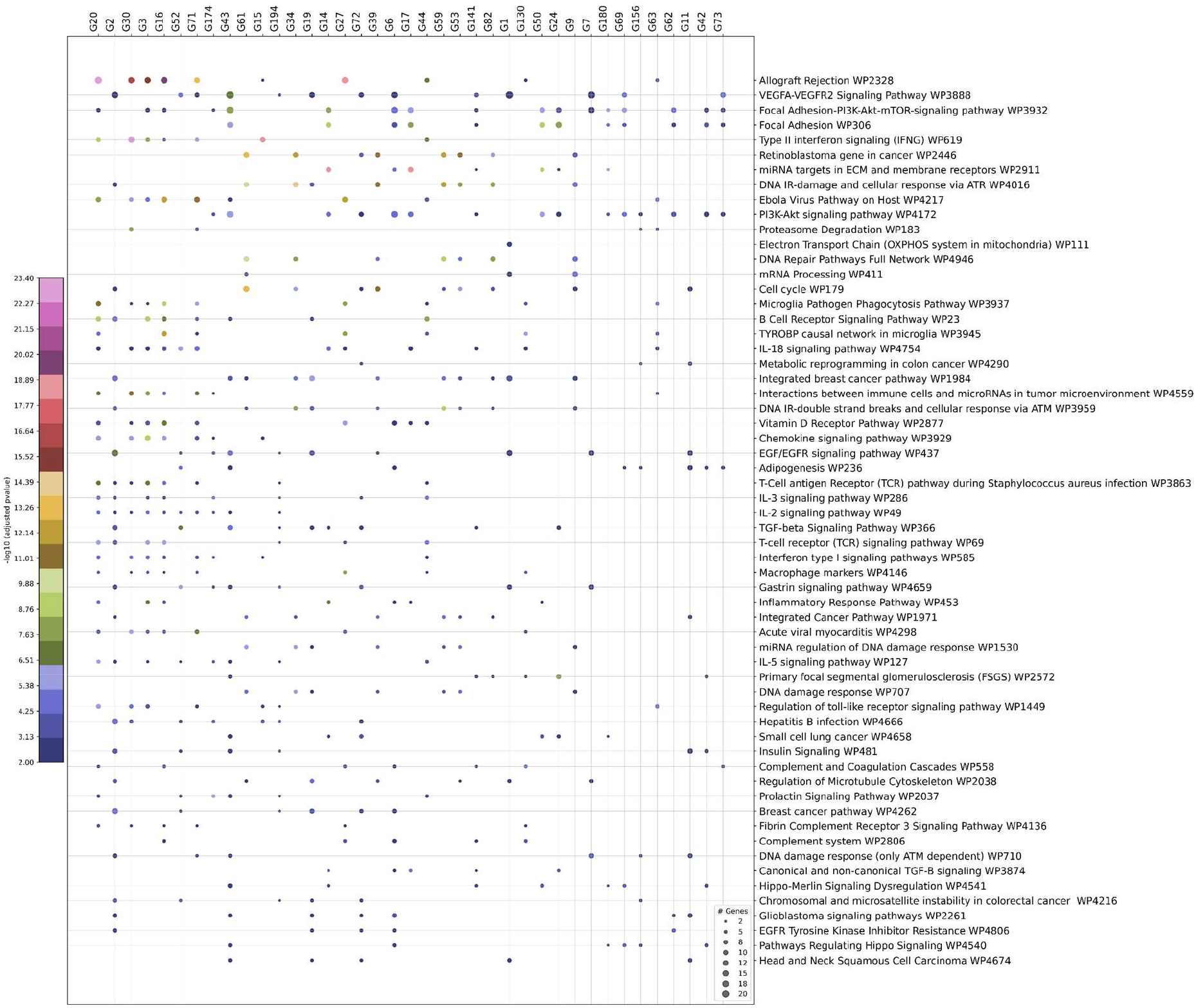
Enrichment of gene groups for cancer WikiPathways is illustrated as 2D scatter plot with the gene group displayed along x-axis and the corresponding enriched pathways on y-axis. The size of the dot represents the number of genes from a specific gene group that has shown enrichment for a particular pathway while its color represents the statistical significance of association in terms of FDR-corrected p-value.

**SFig 6:**
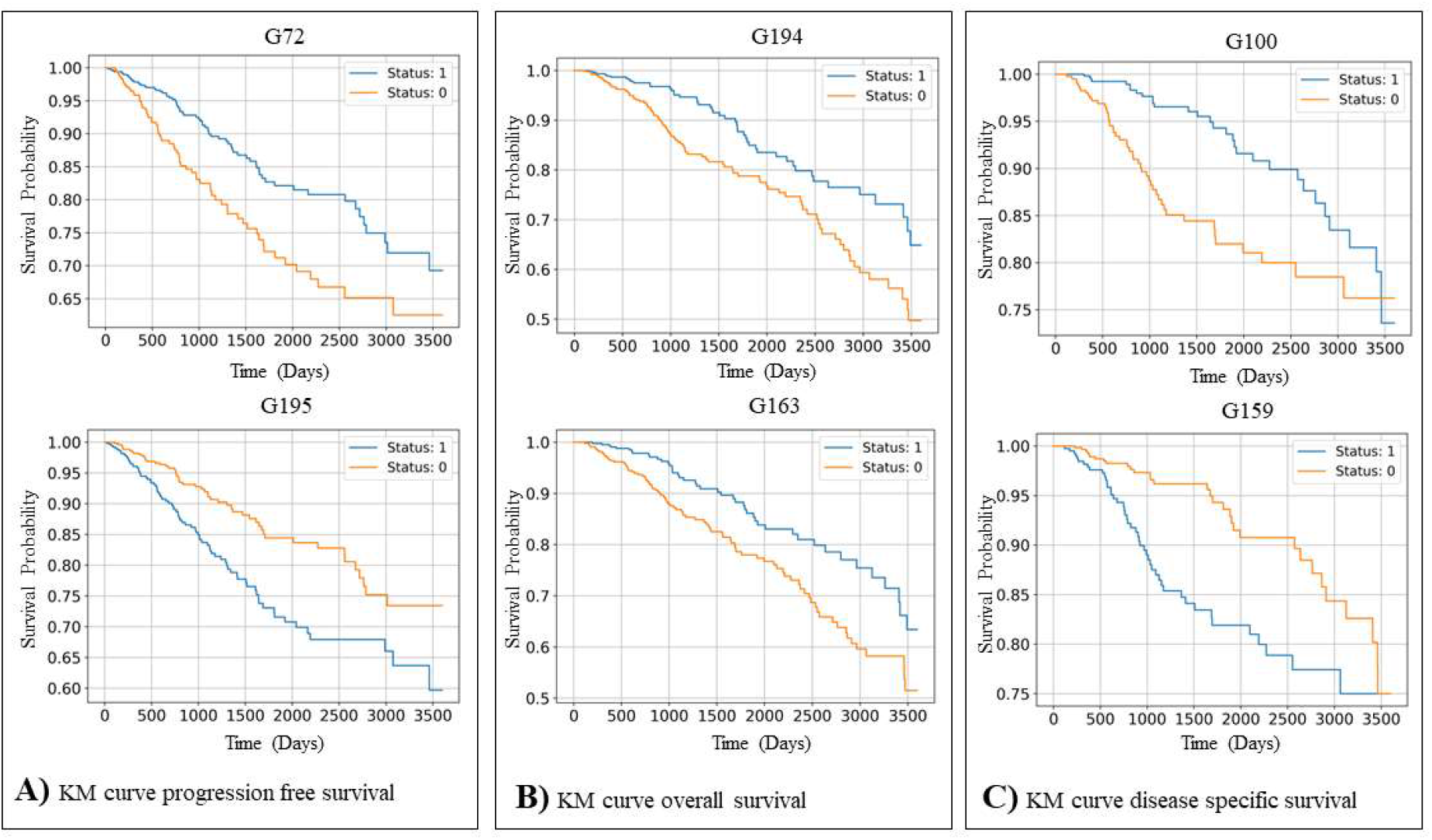
Kaplan-Meier (KM) survival curves of progression-free survival (FPI), overall survival (OS), and disease-specific survival (DSS) of patients stratified based on gene group statuses. A) KM survival curve of PFI of patients based on G72 and G195 status showing that patients can be stratified into high and low risk groups based on G72 and G195 statuses with a significant p-value (log-rank test FDR-corrected p-value < 0.05 as shown in KM survival curve. B) KM overall survival curve of gene groups G194 and G163 are shown. Overall, we found that the binary status of 3 gene groups (G72, G194 and G163) can stratify patients into high and low risk groups with a significant p-value (log-rank test FDR-corrected p-value < 0.05). C) KM-curve of 2 (out of 25) gene groups that shows statistically significant association (log-rank test FDR-corrected p-value < 0.05) with disease-specific survival are shown. Other gene groups that show statistically significant association with DSS are (G72, G163, G194, G80, G123, G82, G10, G76, G165, G156, G120, G53, G142, G25, G144, G61, G113, G150, G151, G175 and G189.

**SFig 7:**
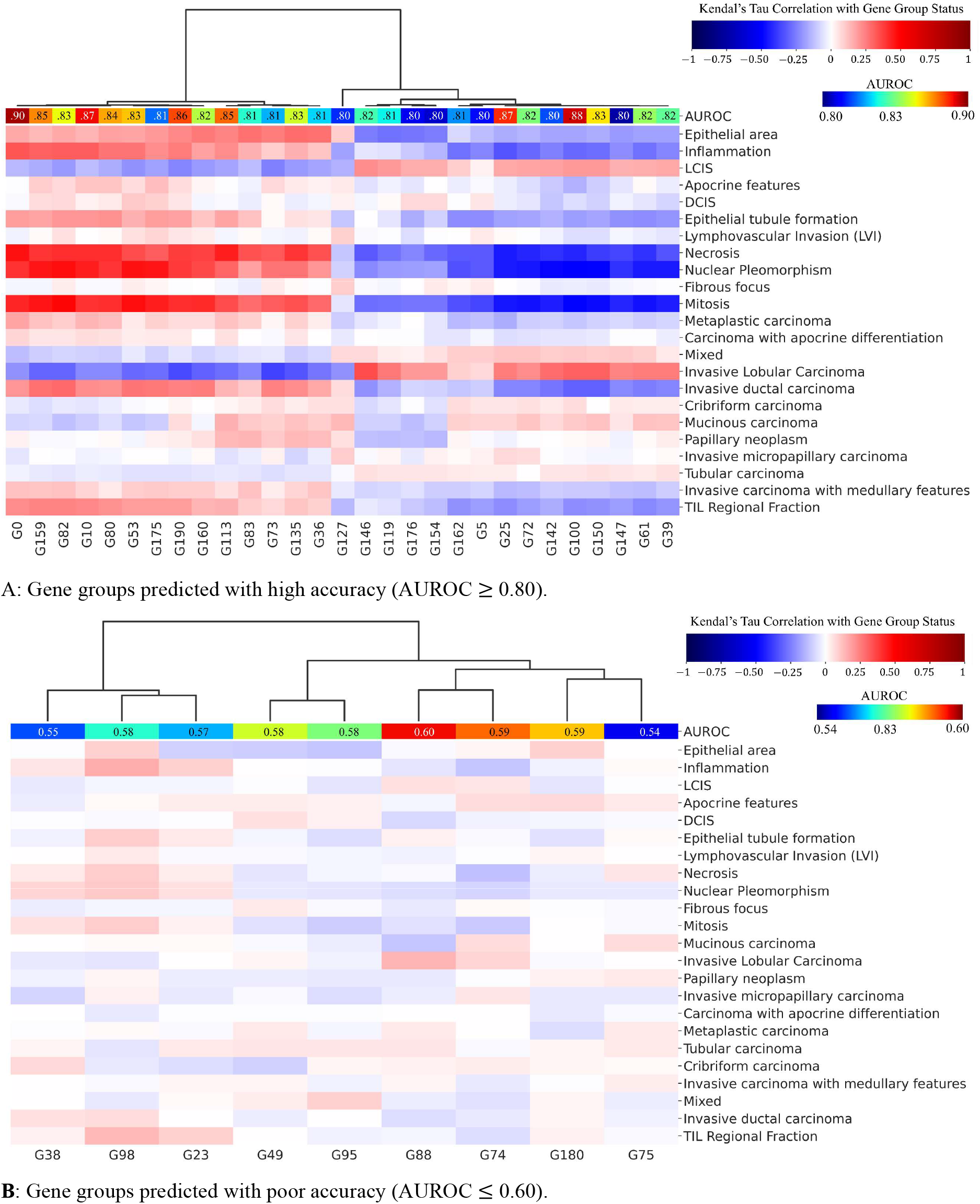
Association of binary statuses of best and worst predicted gene groups with pathologist-assigned histological phenotypes. The plot uses two color bands one for AUROC and one for Kendall’s Tau correlation. The AUROC is illustrated using the jet colormap representing the prediction accuracy of gene group binary status from imaging, while Kendall’s Tau correlation between gene group binary status and various histological phenotypes is shown using the seismic colormap. We also annotated the AUROC colormap with the numeric value representing the mean AUROC value across test folds.

**SFig 8:**
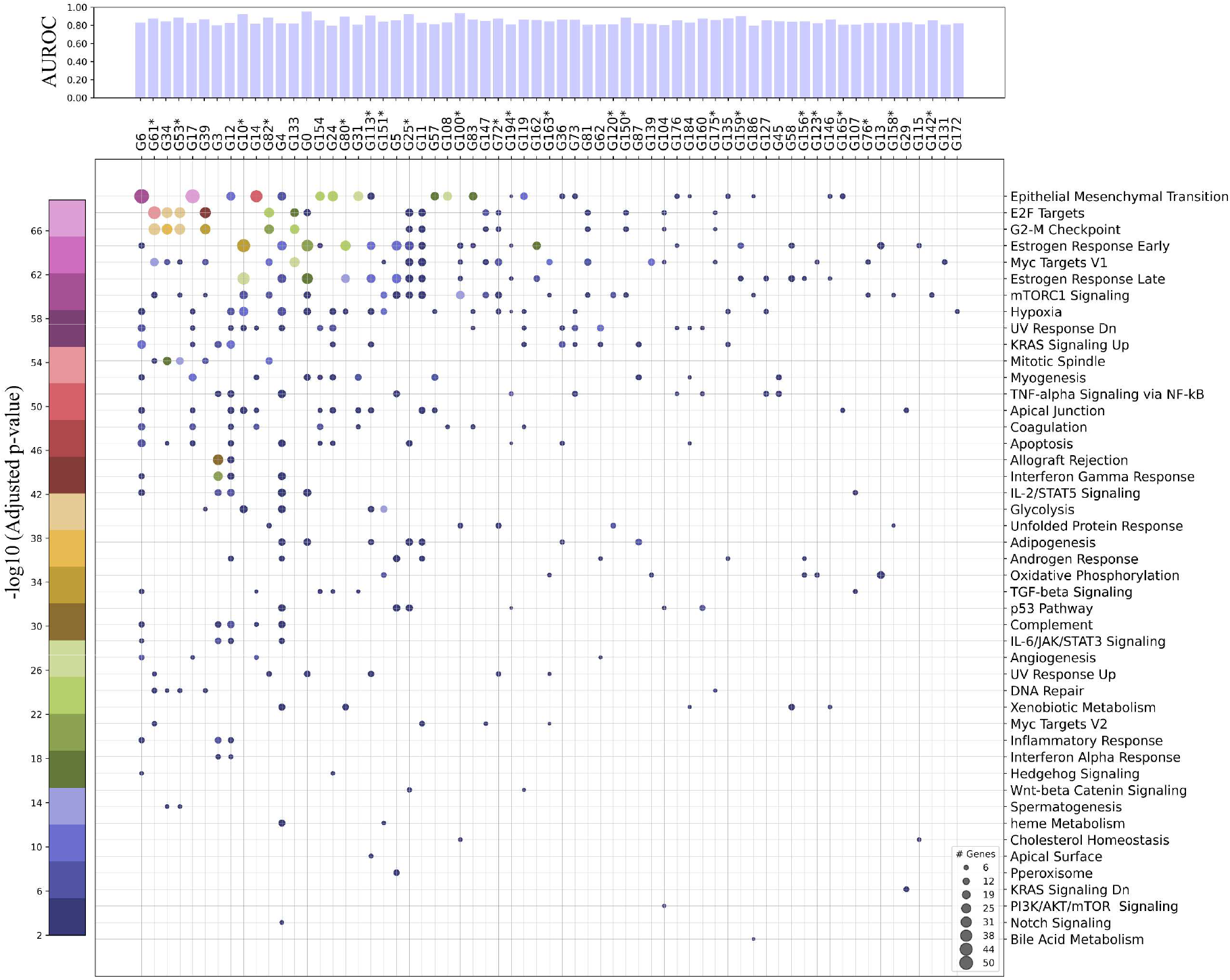
Association of best-predicted gene groups (AUROC ≥ 0.75) with cancer Hallmark processes and disease-specific survival. An example 2D scatter plot showing gene groups (one per column) with hallmark processes (one per row). The size of the scatter dot shows the number of genes in a gene group that has shown statistically significant association (FDR adjusted p-value < 0.01) with a certain biological pathway. In the plot, the p-value is represented by the color of the scatter dots. The top bar plot shows the prediction accuracy (AUROC) at which the binary statuses of these gene groups are predicted from histology images. Furthermore, gene groups that show statistically significant association with disease-specific survival are annotated with a * next to the gene group name.

**SFig 9:**
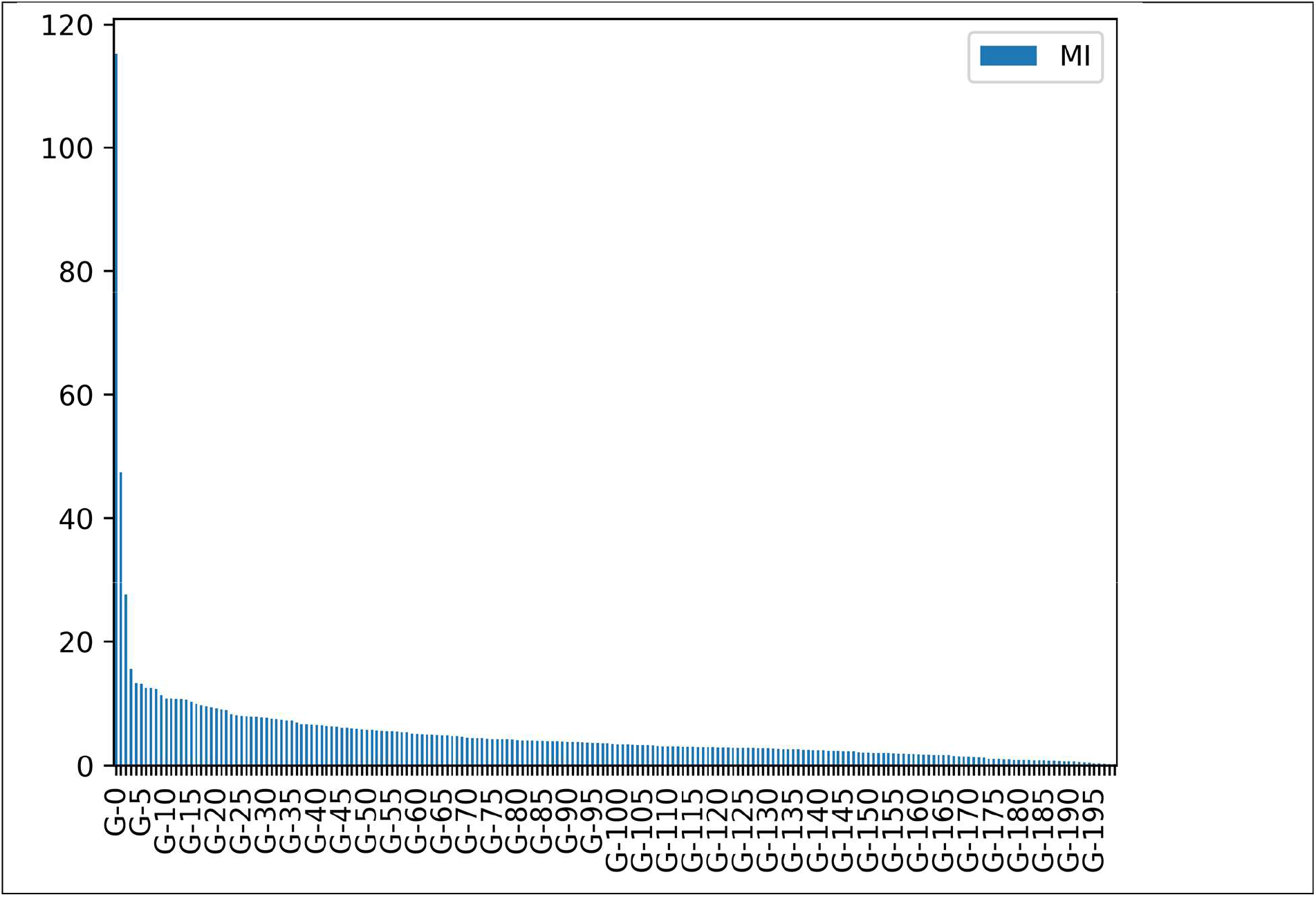
Plot showing the proportion of total correlation explained by each latent factor.

**STable 1:**
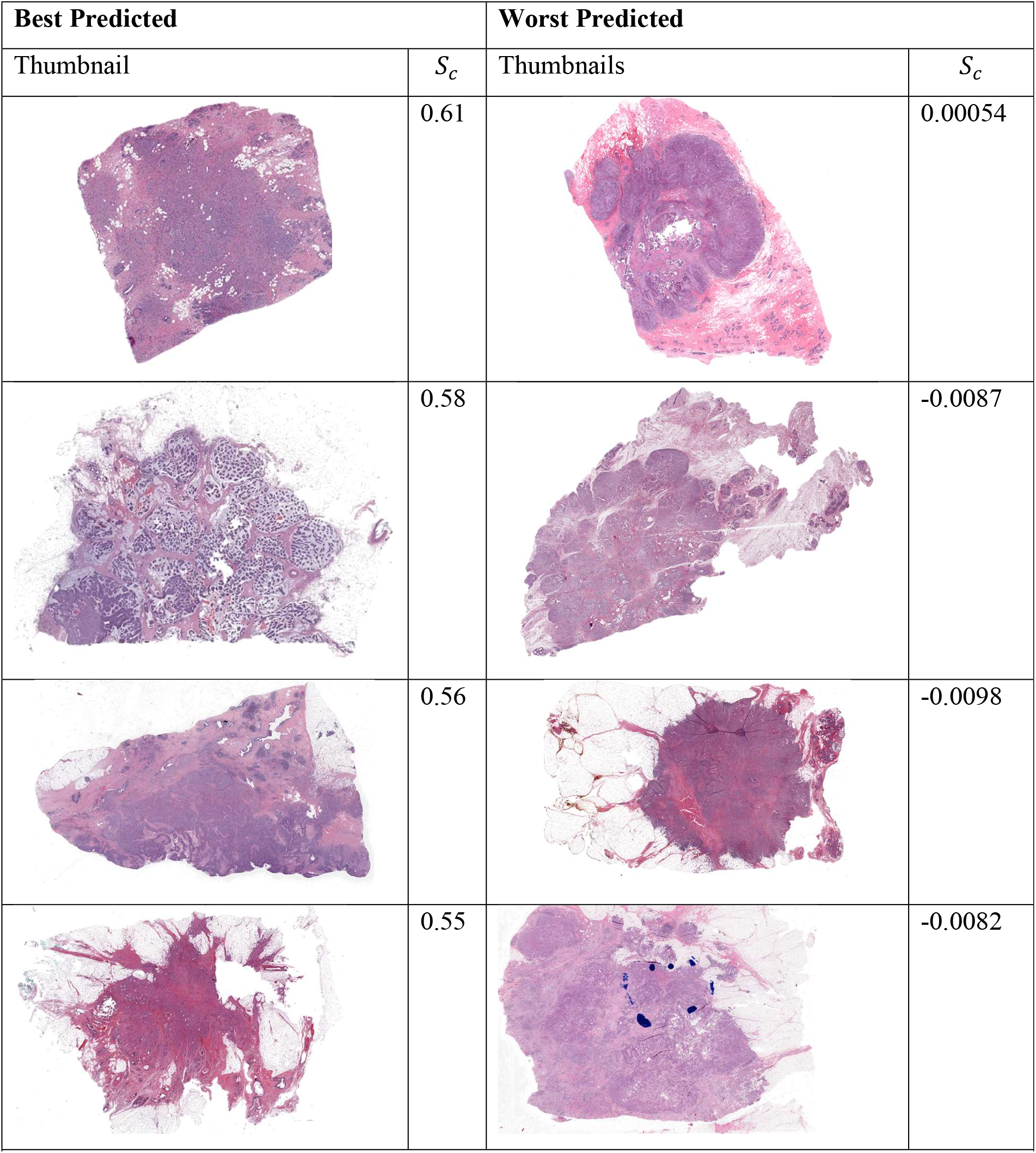
Thumbnails of patient WSIs whose gene expression states are best or poorly predicted from histology images using cosine similarity (𝑆_c_) between ground truth and predicted gene expression states as performance metric.

